# ARPP21 defines a TDP-43-independent aggregation pathway in amyotrophic lateral sclerosis

**DOI:** 10.64898/2026.09.08.750091

**Authors:** Danae Campos-Melo, Allison A. Dilliott, Sali M.K. Farhan, Michael J. Strong, Cristian A. Droppelmann

**Affiliations:** Molecular Medicine Group, Robarts Research Institute, Schulich School of Medicine and Dentistry, Western University, London, ON, Canada; Department of Human Genetics; Department of Neurology and Neurosurgery; The Montreal Neurological Institute and Hospital, McGill University, Montreal, QC, Canada; Department of Pathology and Laboratory Medicine; Department of Clinical Neurological Sciences, Schulich School of Medicine and Dentistry, Western University, London, ON, Canada

**Keywords:** ALS, ARPP21, TDP-43, SZRD1, neurodegeneration, protein aggregates, spinal cord, human iPSC-derived motor neurons

## Abstract

Currently, pathological inclusions of ubiquitinated TDP-43 are considered central to the pathogenesis of amyotrophic lateral sclerosis (ALS). However, this view has yielded sparse attention to covert alternative pathways of aggregation in the disease. Here, we identified a pathological axis independent of TDP-43 led by ARPP21, a SUZ domain-containing RNA-binding protein encoded by a gene recently described in strong association with ALS. ARPP21 showed large fibrillar non-ubiquitinated aggregates in all the non-SOD1 ALS cases studied, consistently segregating from TDP-43 pathology. Biophysically, ARPP21 undergoes spontaneous phase separation, with a higher propensity for condensate formation than TDP-43 but without co-aggregating with it. ARPP21 also exhibited very slow soluble behavior and lower condensate dynamics compared to TDP-43, an effect that was more pronounced for the ARPP21-P529L disease-related variant. In human iPSC-derived motor neurons, expression of *ARPP21* variants was sufficient to drive condensate formation and reduce cell viability. Mechanistically, ARPP21 promoted intercellular propagation by inducing tunnelling nanotube formation, enabling efficient cell-to-cell spreading of aggregates. Genetic analyses further identified multiple *ARPP21* variants in ALS, supporting its clinical relevance. Together, our findings uncover ARPP21 as a previously unrecognized TDP-43-independent aggregation pathway in ALS, with implications for disease heterogeneity and therapeutic targeting.

## Introduction

Since the identification of TDP-43 as a component of ubiquitinated aggregates in ALS and FTD approximately 20 years ago (*1, 2*), the majority of ALS research has been focused on understanding the complexity of the multilayer role of TDP-43 in the pathology (*3–5*). The presence of TDP-43 pathology in ∼97% of all ALS cases, which includes TDP-43 cytoplasmic mislocalization and inclusion formation, has fostered these studies even though pathogenic mutations in the gene that encodes TDP-43, *TARDBP*, occur in < 1% of ALS cases (*6–9*).

After 2006, the presence of TDP-43 positive ubiquitinated aggregates in tissue samples as a routine histopathological inspection in ALS contributed to this TDP-43 centric approach, often leading us to ignore the possibility of ubiquitinated or non-ubiquitinated TDP-43 negative aggregates that might have a role in the pathology of ALS.

Recently, mutations in the cAMP-regulated phosphoprotein 21 gene (*ARPP21*), a member of a poorly characterized family of RNA-binding proteins (RBPs), were described in small cohorts of ALS patients from the UK, Spain, and China (*10–12*). This prompted us to investigate the pathological and spreading characteristics of ARPP21. While this manuscript was in preparation, a large genetic study reported a robust association between the *ARPP21* gene and ALS (*13*).

Here, we show that ARPP21 forms large fibrillar aggregates in ALS tissue that are independent of the TDP-43 pathological state. ARPP21 aggregates are present regardless of the ALS type and mutation, with the exception of mtSOD1 samples, and are non-ubiquitinated in nature. In cultured cells, ARPP21 condensates are easily induced, showing a complete segregation from TDP-43, but often co-aggregating with SUZ domain-containing protein 1 (SZRD1), another member of the small group of SUZ domain-containing RBPs. Fluorescence recovery after photobleaching (FRAP) showed that ARPP21condensates are less dynamic than those of TDP- 43, an effect that was more pronounced in the ALS-associated ARPP21-P529L variant. Soluble ARPP21 studied by fluorescence correlation spectroscopy (FCS) identified a slow oligomeric ARPP21 component in the cytoplasm. In human iPSC-derived motor neurons, we observed that the transduction with AAV9/ARPP21-wt and AAV9/ARPP21-P529L led to condensate formation and decreased cell viability. Moreover, ARPP21expression induced the formation of tunnelling nanotubes (TNTs), suggesting an efficient spreading mechanism between cells. Finally, a curated genetic analysis of human databases showed several *ARPP21* variants in ALS compared to controls, strengthening the importance of this gene in ALS studies.

## Methods

### Tissue samples

Human spinal cord tissues from 5 neurologically normal controls and 14 ALS patients were extracted from the Strong lab ALS/MND tissue bank (Supplementary Table 1, patient demographics). All tissues were obtained using ethics research protocols approved by the Western University Health Sciences Research Ethics Board (HSREB) (HSREB #103735 & #124855).

### Immunohistochemistry

7mm- tissue FFPE tissue slices were warmed at 60°C for 30 min and rehydrated in a series of graded alcohols and water. Antigen retrieval was performed in a pressure cooker for 30 min at 100°C in a buffer containing 10 mM sodium citrate and 0.05% Tween, pH 6.0. Then, slides were blocked for 60 min at room temperature in PBS pH 7.2 solution with 5% BSA and 0.3% Triton-X-100, and incubated with primary antibodies at 4°C overnight in a humidifying chamber. After, slides were incubated with Alexa Fluor secondary antibodies for 60 min at room temperature.

Nuclear staining was performed using 2 μg/ml Hoechst for 3 min. Primary and secondary antibodies and working dilutions are indicated in Supplementary Table 2. Validation of ARPP21 antibody is shown in Supplementary Figure 1A and B.

### Constructs, RT-PCR and sequencing

The coding regions of the ARPP21 spinal cord isoforms (ARPP21^sc^-wt, ARPP21^sc^-P529L, ARPP21^sc^-E377K) and SZRD1 were synthesized with a myc tag and cloned into the pcDNA3.1 vector (GeneScript, USA) for mammalian cell expression. The viral sequences containing ITRs, CMV enhancer, *SYN1* promoter, WPRE element, SV40 polyA signal and coding regions of EGFP, ARPP21^sc^-wt or ARPP21^sc^-P529L were synthesized and cloned into pUC18 (GeneScript, USA) for AAV9 generation (VectorBiolabs, USA). The coding regions of TDP-43-wt, ARPP21^sc^-wt, ARPP21^sc^-P529L, ARPP21^sc^-E377K were subcloned into the pHA-Clover vector for mammalian cell expression. For the detection of the ARPP21 isoforms, total RNA was extracted from three healthy spinal cord tissue samples, and then RT-PCR was performed using primers that allowed us to detect the presence or absence of the exons 9, 11 and 15 in the mature ARPP21 mRNA. For the sequencing, spinal cord DNA from all the tissues used for pathology were extracted, then intronic/exonic specific regions surrounding exons 13 and 15 of the *ARPP21* gene were amplified by PCR. The PCR products were purified and then sequenced (Sanger method) using the same primers to discard the presence of ALS-linked *ARPP21* mutations P529L and E377K. The lists of critical primers and the plasmids utilized in this study are shown in Supplementary Table 3 and Supplementary Table 4, respectively.

### Cell culture and transfections

HEK293T and SH-SY5Y cells were cultured in Dulbecco’s Modified Eagle’s Media (DMEM) containing 10% Fetal Bovine Serum (FBS), at 37C° with 5% CO_2_. Transfections with ARPP21 constructs (Supplementary Table 4) in 6-well plates (HEK293T) and 96-well plates (SH-SY5Y) were performed using Lipofectamine 2000, following the manufacturer’s instructions. For stress, cells were incubated with 500mM NaAsO4 (sodium arsenite) for 1h.

### Immunofluorescence and quantification

Cultured cells were washed and fixed with 4% PFA for 15 minutes at room temperature, followed by permeabilization with 0.2% Triton X-100 and blocking with 4% BSA. Then, cells were incubated with primary antibodies overnight at 4° C and the next day with fluorescent secondary antibodies for 1h at room temperature (Supplementary Table 2). Samples were stained for nuclei using Hoechst and mounted using fluorescent mounting media (Dako). Imaging was accomplished with a Leica TCS SP8 confocal microscope. Fiji was used for quantification of condensates and TNTs per cell.

### Co-localization images

Intensity Correlation Analysis (*14*) using ImageJ software was performed to obtain the co- localization images. Co-localized pixels are shown as PDM (Product of the Differences from the Mean) images [PDM = (red intensity-mean red intensity)×(green intensity-mean green intensity)]. In the images, a scale showing the degree of colocalization was added (where blue and purple colors indicate a lower level of co-localization, while yellow and white indicate a high level of co-localization).

### Protein modelling

Prediction and visualization of the structures of ARPP21^sc^-wt and P529L and E377K variants, and SZRD1, were performed using AlphaFold Server (*15*) and PyMOL.

### Fluorescence recovery after photobleaching (FRAP)

FRAP experiments were performed using ARPP21^sc^-wt, ARPP21^sc^-P529L, or ARPP21^sc^-E377K fused to clover (Figure 5A) or TDP-43-wt-clover in HEK293T cells in DMEM/HEPES culture medium (ThermoFisher) 48 hours after the transfection using a Leica TCS SP8 confocal microscope with a 40x objective lens and the LAS X software. For photobleaching, we excited a 1 µm^2^ region-of-interest (ROI) with a 488 nm diode laser at 100% power for 0.5 s. Clover fluorescence recovery after bleaching was monitored for 1 min. For each clover fusion, 14 aggregates were measured from 3 independent experiments. Curves were fitted using the ‘one-phase association’ equation in GraphPad Prism 11. In this equation computed values “span” indicates the total change in Y (plateau – Y_0_); “tau (τ)” is the time constant and indicates the amount of time necessary to reach 63.2% of the way from Y_0_ to the plateau; and, “k” is the rate constant, expressed in reciprocal units (s^-1^), which indicates how quickly the system approaches the plateau.

### Fluorescence correlation spectroscopy (FCS)

HEK293T cells cultured in live cell imaging dishes were transiently transfected with clover-tagged ARPP21sc-wt, ARPP21sc-P529L, and ARPP21sc-E377K variants, TDP-43-wt-clover or clover alone (Figure 5A). Experiments were performed in regions of the cells that did not show condensate formation 24 hours after transfection to decrease the chance of finding high number of spontaneous condensates. Fluorescence microscopy measurements were performed in a Leica SP8 confocal microscope using a 63x objective and a hybrid detector operating in photon counting mode. Data were acquired for Alexa Fluor 488 Azide solution and clover proteins from transfected cells using XT line scan and X-segmentation, each 8192-pixel line divided into 8 segments of 1024 pixels (fast molecules) or XT line scan and Y-segmentation in which each 204,800 pixels column was divided into 32 segments of 6,400 pixels for ARPP21sc-wt-clover, ARPP21sc-P529L-clover, ARPP21sc-E377K-clover, and TDP-43-wt-clover (slower molecules). Images were opened and processed using a previously described custom MATLAB script developed specifically for this type of analysis (*16*). Autocorrelation (ACF) versus time curves fitting was run in GraphPad Prism 11.

Stable fits of the ACF curves were obtained with a one-component lateral anomalous diffusion model using equation (1):

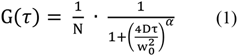

Where G(t) is the autocorrelation function (Y), N is the number of fluorescent particles in the focal volume, τ (tau) is the scan time (X), w_o_ is the lateral radius or beam waist, D is the diffusion coefficient, and α is the anomalous component (*16, 17*). Fitting quality was evaluated by visual inspection, stability of the parameters in the replicates, R-squared values, and residual analysis. W_0_ for the instrument was obtained by calibrating an ACF curve of a 10 nM solution of Alexa Fluor 488 Azide in PBS and fitting with equation (1) using a value of α = 1. As the diffusion coefficient of Alexa Fluor 488 is known (D_Alexa 488_ = 395µm^2^/s), the w_0_ for our instrument was 0.2745 (Supplementary Figure 1C).

Molecular brightness was calculated with:

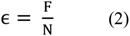

where ∈ is brightness, F is average fluorescence during acquisition, and N is the number of fluorescent particles in the focal volume (*18, 19*). Values were normalized to the monomeric control (clover).

### Human iPSC-derived motor neurons

Human iPSC-derived motor neurons from spinal cord (iXCells Biotechnologies) obtained from healthy individuals were seeded at a density of 10^5^ cells/cm^2^ in CC2 8-well chamber slides (Thermo Fisher; immunocytochemistry) or 96-well plates coated with Matrigel (viability assays) and maintained at 37 C° with 5% CO_2_ in a humidified chamber in motor neuron culture medium kit (iXCells Biotechnologies), following the manufacturer’s instructions. Cell morphology was monitored to avoid large clumps, which was optimal after 7 days of culture.

### AAVs transduction

Using the pAAV-GFP, pAAV-ARPP21^wt^, and pAAV-ARPP21^P529L^ plasmids, the adeno-associated viruses serotype 9 (AAV9) for neuronal-specific expression of GFP (AAV9-GFP), ARPP21^wt^ (AAV9- ARPP21^wt^), and ARPP21^P529L^ (AAV9- ARPP21^P529L^) were produced to a yield of 1.38x10^12^ GC/ml, 1.7x10^12^ GC/ml, and 1.57x10^12^ GC/ml, respectively (Vector Biolabs). Cells were transduced to a 1x10^4^:1 GC:cell ratio. After 72 hours, cells were analyzed for imaging or viability assay.

### ATP viability assay

Cells were seeded in 96-well plates at 10^4^ cells/well for SH-SY5Y or 10^5^ cells/cm^2^ for human iPSC-derived motor neurons. SH-SY5Y cells were transfected with ARPP21 constructs using Lipofectamine 2000 while human iPSC-derived motor neurons were transduced 7 days post-seeding using AAV9 containing the coding region for GFP, ARPP21^sc^-wt or ARPP21^sc^-P529L. Cell viability was quantified using CellTiter-Glo® 2.0 Assay (Promega) following the manufacturer’s instructions. After transfection of the ARPP21 constructs (SH-SY5Y) or transduction of AAV9 viruses (Supplementary Table 4) containing ARPP21 WT or mutant (iPSC-derived MNs), the CellTiter-Glo® 2.0 reagent, was added and luminescence was measured in a luminometer.

### Variant genomic analysis

To assess for rare *ARPP21* variants that may be associated with ALS, we interrogated the ALS Knowledge Portal, now referred to as the Neurodegenerative Disease Knowledge Portal (*20, 21*) (individuals with ALS = 3864, controls = 7839) quality-controlled exomes and Project MinE ALS Consortium (*22*) (individuals with ALS = 6596, controls = 2454) genome dataset for variants within coding regions (± 10 nucleotides of intron-exon junctions) of the gene. Variants were annotated using the Ensembl Variant Effect Predictor (GRCh 37, NM_001385562.1; accessed July 2025), including variant consequence, gnomAD allele frequencies (exome and genome), AlphaMissense (*23*) predicted pathogenicity classification, and SpliceAI (*24*) predicted splicing effects. Only rare variants were retained, specifically those with an allele frequency < 0.01 or those not found in the gnomAD exome and genome datasets. Variants were then binned into four types: 1) synonymous variants; 2) missense variants, including missense single nucleotide variants (SNVs) and in-frame insertions/deletions; 3) damaging missense variants, defined as those predicted “pathogenic” using AlphaMissense; and 4) putative protein truncating variants (PTVs), including frameshift, splice acceptor, splice donor, and stop gain variants. Figures were made using the St. Jude PeCan Data Portal ProteinPaint (*25*) tool.

### Statistical analysis

The statistical analyses were performed with GraphPad Prism 11 software. One-way ANOVA with Tukey’s post-hoc test were performed. Data were expressed as mean ± SEM and judged to be statistically significant when p < 0.05.

## Results

### ARPP21 forms non-ubiquitinated fibrillar aggregates in the ALS spinal cord

After *ARPP21* genetic variants were reported in small cohorts of ALS patients from the UK, Spain, China (*10–12*), and more recently in France (*26*), we decided to study ARPP21 pathology in spinal cord tissues from ALS patients. We performed ARPP21 immunohistochemistry of spinal cords of 14 ALS patients and 5 controls. The sequencing of spinal cord genomic DNA from the patients used in this study did not show the *ARPP21* variants P529L and E377K previously described (*10–12*).

Tissue exploration showed low ARPP21 expression in small nuclear and cytosolic granules in control spinal motor neurons (Figure 1A and Supplementary Figure 2A), and rare cells showing larger ARPP21 granules (Supplementary Figure 2B). In all sporadic and familial ALS cases studied (n=11) with the exception of I113T and A4T mtSOD1 cases, the presence of fibrillar ARPP21 aggregates reminiscent of crystal-like structures was observed in the cytoplasm of motor neurons (Figures 1 and 2; Supplementary Figure 2). Co-immunostaining with TDP-43 and 3D analysis showed no colocalization between ARPP21 and TDP-43 when aggregates for both RBPs were present, even in the same cell (Figure 1A and B). Interestingly, no trace of TDP-43 pathology was observed in most motor neurons when ARPP21 cytoplasmic aggregates were present. Even an ALS case that lacked TDP-43 pathology showed large ARPP21 aggregates (Figure 2A; Supplementary Figure 2A). mtSOD1 cases showed consistent granular ARPP21 distribution (Figure 2A) similar to controls. ARPP21 aggregates in ALS showed no ubiquitin staining (Figure 2B, C), even though ubiquitin aggregates were clearly observed in neurons with no ARPP21 pathology in the same tissue (Figure 2C). This observation was confirmed in the case negative for TDP-43 pathology (Supplementary Figure 2C).

**Figure 1.**
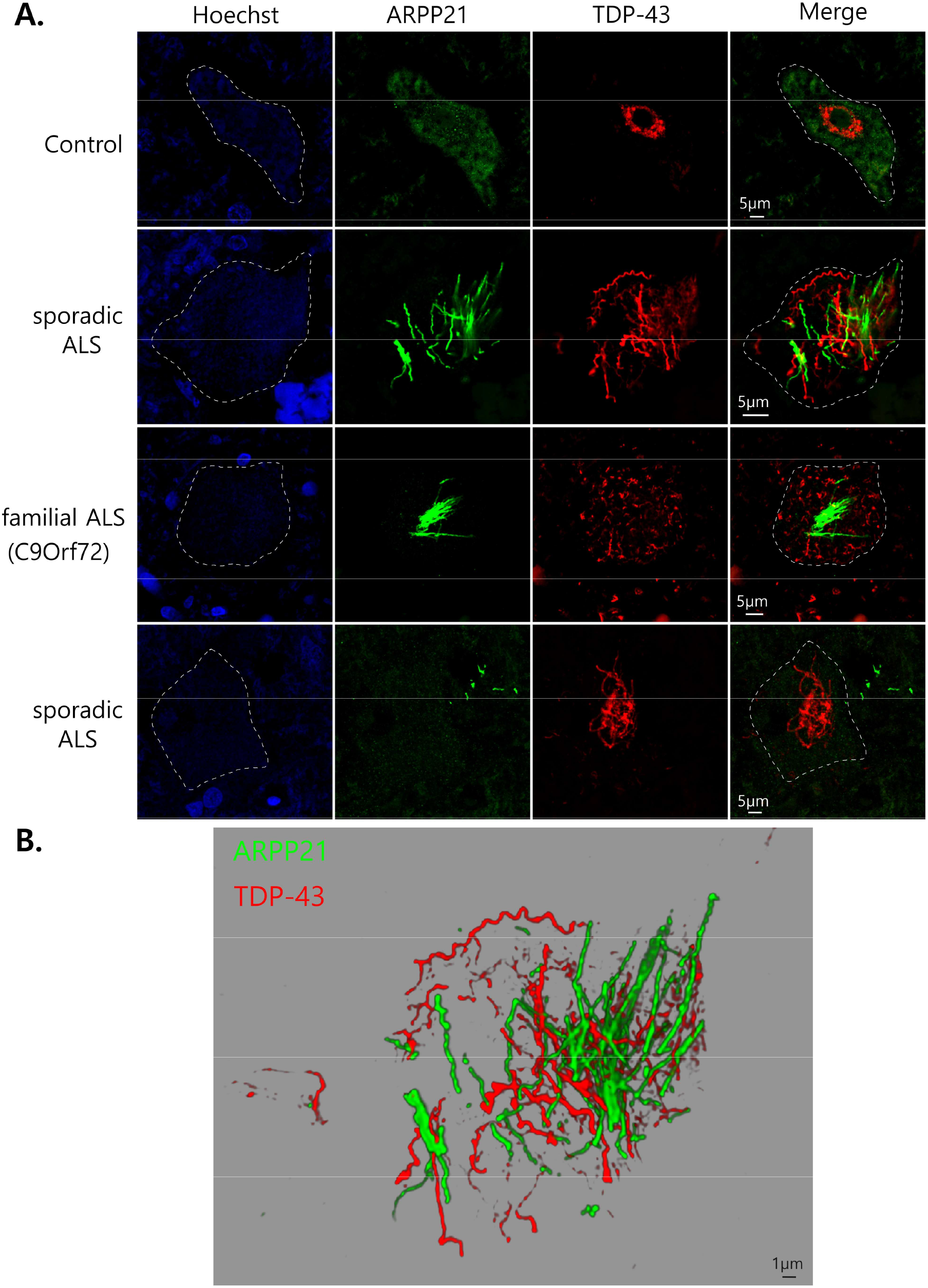
**ARPP21 forms fibrillar aggregates that do not co-localize with TDP-43 in the spinal cord in sporadic and familial ALS**. (**A**) ARPP21 (green) shows mostly granular localization in the cytoplasm of motor neurons in the control spinal cord (top panel). In sporadic ALS and a familial C9Orf72 ALS case, ARPP21 forms fibrillar aggregates in motor neurons that do not co-localize with TDP-43 aggregates (red; middle panels) or could be present in neighbor cells (sporadic ALS, bottom panel). Two different sporadic ALS cases are shown. A segmented white line demarcates the borders of motor neurons. (**B**) 3D reconstruction of large ARPP21 and TDP-43 motor neuron aggregates (sporadic ALS, second panel from top) demonstrates the lack of co-aggregation between ARPP21 and TDP-43.

**Figure 2.**
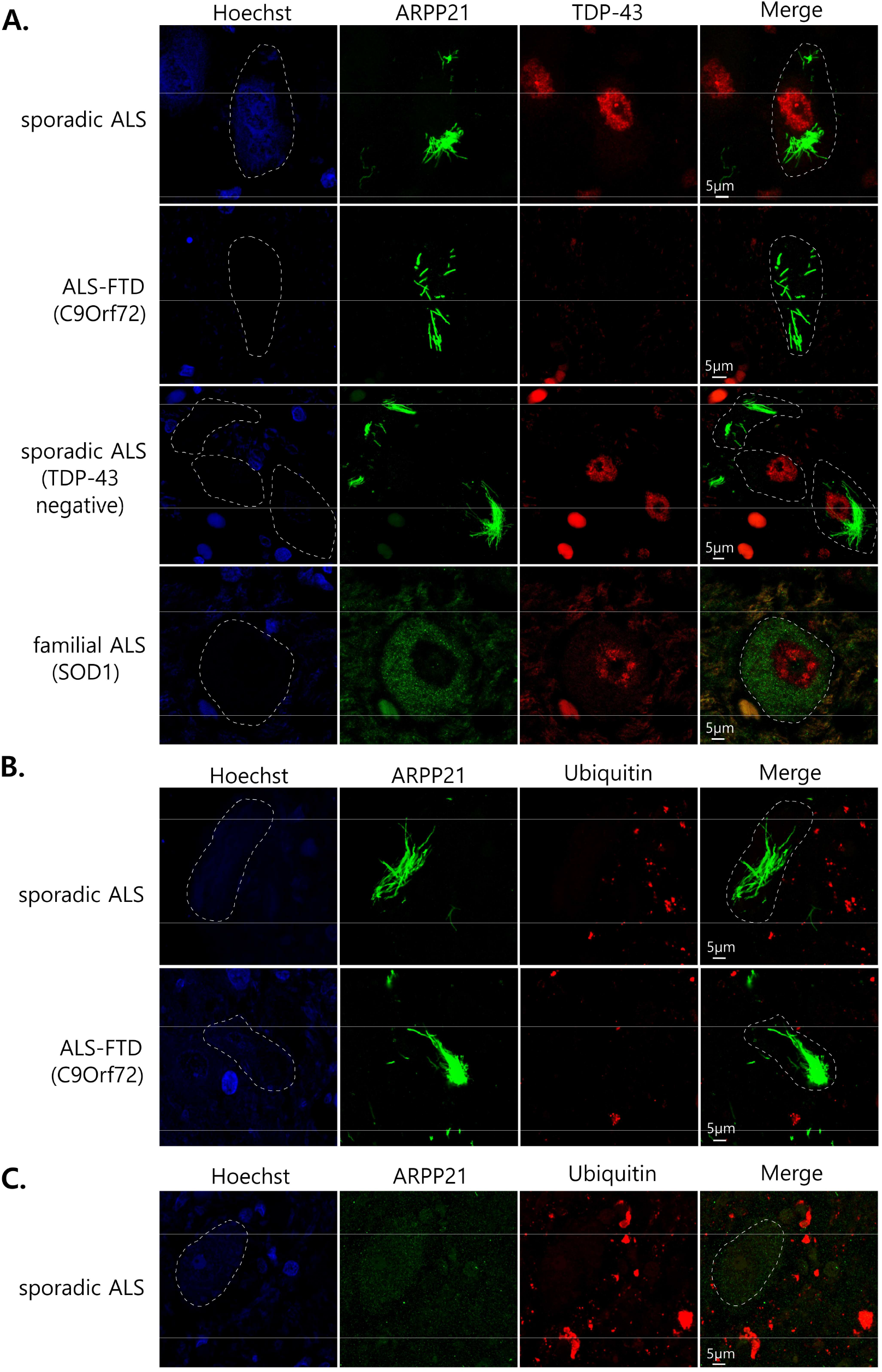
ARPP21 aggregates are present in motor neurons without TDP- 43 pathology in different ALS cases and are non-ubiquitinated. (**A**) sporadic ALS and familial ALS (C9Orf72) cases show ARPP21 aggregates without TDP-43 pathology in the same motor neurons (top panel and second panel from the top). A case that was reported negative for TDP-43 pathology (third panel from top) shows large ARPP21 aggregates. mtSOD1 cases analyzed do not present ARPP21 pathology (bottom panel). (**B**) ARPP21 aggregates are not ubiquitinated in the ALS patients analyzed, but large ubiquitinated ARPP21- negative aggregates are observed in the same tissues (**C**). A segmented white line demarcates the borders of neurons.

In sum, these results indicate that ARPP21 forms fibrillar aggregates that are independent of TDP-43 pathology in the ALS spinal cord.

### ARPP21 is prone to form condensates in cells

To study ARPP21 in cultured cells, we first evaluated splicing variants expressed in the human spinal cord. Previously, four different isoforms of *ARPP21*, including the thymocyte specific ARPP21-related called *TARPP* (*27*), were reported (Supplementary Figure 3A; Accession numbers: NM_001385592; NM_001385486; NM_001267617; NM_016300). Using a specific set of primers (Supplementary Figure 3B; Supplementary Table 3), we detected three ARPP21 isoforms in the spinal cord: ARPP21^847aa^, ARPP21^813aa^ and ARPP21^793aa^. From this group, ARPP21^813aa^, which lacks exon 9, was the most abundantly expressed. We did not detect the *TARPP* (ARPP21^812aa^) isoform (Supplementary Figure 3B and C). We cloned the myc-tagged ARPP21^813aa^ -wt, and P529L and E377K variants previously described in ALS patients (*11, 12*) (P563L and E411K, respectively in the full-length ARPP21^847aa^ isoform), here after called ARPP21^sc^-wt, ARPP21^sc^-P529L, and ARPP21^sc^-E377K for <u>s</u>pinal <u>c</u>ord isoforms (Figure 3A), and we expressed these constructs in HEK293T cells.

**Figure 3.**
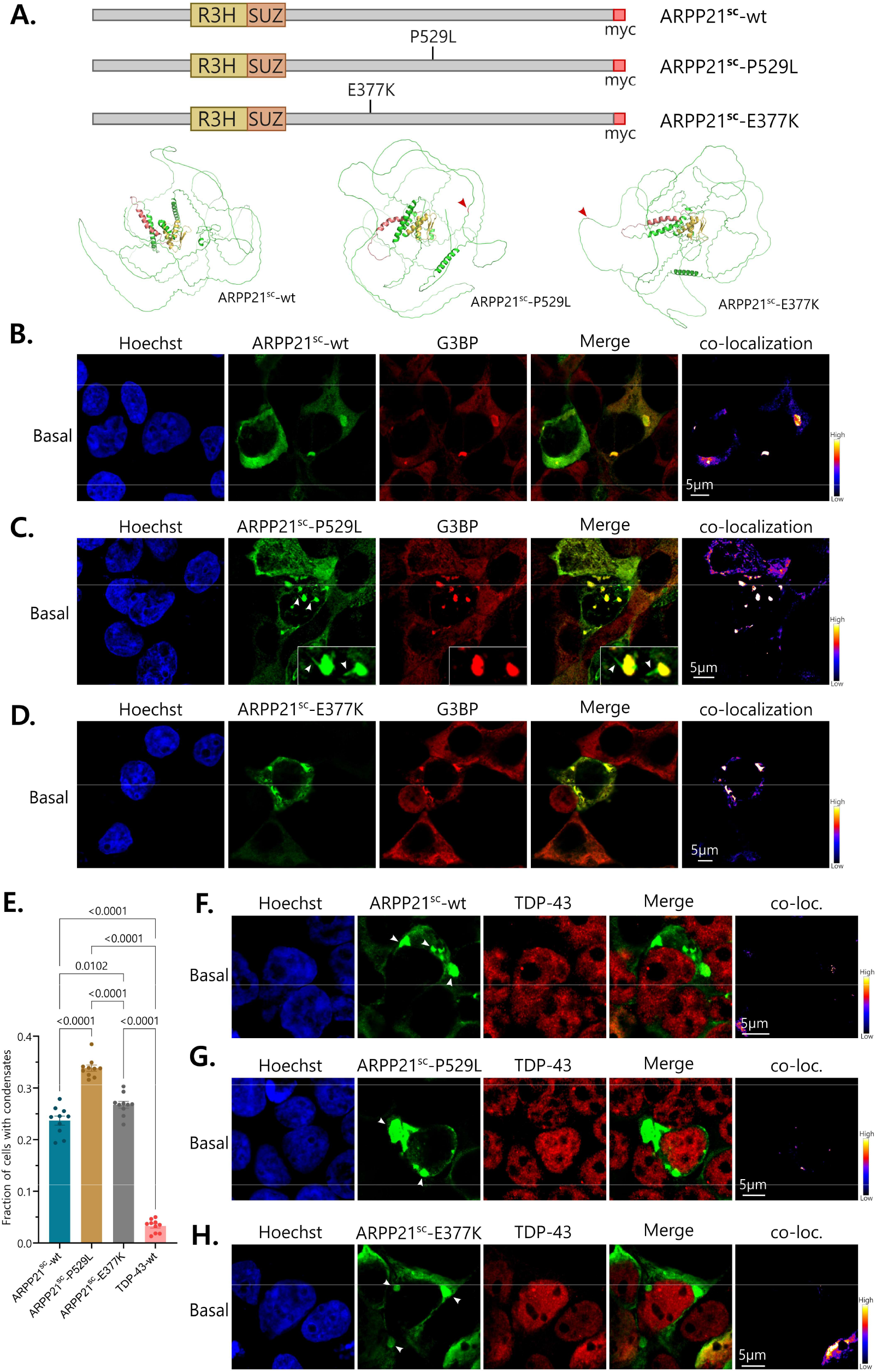
ARPP21 is prone to aggregation in cells. (**A**, top) Schematic showing the myc-tagged spinal cord (sc) isoform of ARPP21-wt, and P529L and E377K variants. (**A**, bottom) Alphafold modelling of ARPP21^sc^-wt and the variants studied. The position of the P529L and E377K mutations is indicated by red arrowheads. ARPP21 is 87% disordered (MobiDB). R3H (yellow) and SUZ (red) domains are highlighted in the structures. (**B, C, D**) ARPP21^sc^-wt, and P529L and E377K variants form spontaneous condensates that co- localize with G3BP under basal conditions in HEK293T cells. White arrowheads in C indicate ARPP21-positive G3BP-negative fibrils emerging from condensates (magnified in insets). (**E**) Quantification of the percentage of cells with ARPP21^sc^ and TDP-43-wt spontaneous condensates under basal conditions. ARPP21^sc^-P529L forms significantly more condensates than ARPP21^sc^-wt and ARPP21^sc^-E377K. All ARPP21^sc^ variants show more than 7-fold condensates than TDP-43-wt. (**F, G, H**) ARPP21^sc^-wt and variants condensates (white arrowheads) do not co-aggregate with TDP-43 in basal conditions.

AlphaFold modelling showed the highly disordered predicted structure of ARPP21 (87.3% disordered compared to 60.6% value for TDP-43 using MobiDB (*28*); https://mobidb.org) and the positions of P529L and E377K mutations in the disordered regions (Figure 3A). Under basal conditions, the expression of ARPP21^sc^-wt, ARPP21^sc^-P529L, and ARPP21^sc^-E377K variants easily induced the formation of spontaneous cytoplasmic condensates (Figure 3B, C and D).

Aggregate quantification under basal conditions showed a higher number of cells with ARPP21^sc^- positive condensates when P529L or E377K variants were transfected compared to the wt (Figure 3E). P529L was more prone to induce condensates in cells than E377K. ARPP21^sc^-wt, ARPP21^sc^-P529L, and ARPP21^sc^-E377K showed 7.2, 10.3, and 8.1-fold more condensates than TDP-43-wt under basal conditions, respectively. ARPP21^sc^-positive condensates always co- localized with the stress granule marker G3BP, which suggests the formation of aberrant condensates (Figure 3B, C and D). Short fibrillar structures of ARPP21^sc^ that are G3BP-negative were occasionally observed emerging from droplet-like condensates (Figure 3C, insets). After oxidative stress using NaAsO_2_, we observed an increase in G3BP-positive condensates, the majority of which were ARPP21^sc^-positive, both for wt and P529L and E377K variants (Supplementary Figure 4A, B, C and D). Co-immunostaining with TDP-43 showed ARPP21^sc^- positive aggregates almost completely segregated from TDP-43 under basal and stress conditions. The expression of the ARPP21^sc^ variants did not induce endogenous TDP-43 condensate formation or nuclear clearance, which supports the idea of independence between ARPP21 and TDP-43 pathology (Figure 3F, G and H, and Supplementary Figure 4E, F and G).

Next, we did immunostaining in human tissue for SZRD1, another member of the SUZ family of four RBPs. We noticed, that unlike ARPP21, SZRD1 forms different types of aggregates in the spinal cord in ALS, although not often in motor neurons, and that these aggregates show co-aggregation with ARPP21 to a variable extent (Figure 4A and Supplementary Figure 5A).

**Figure 4.**
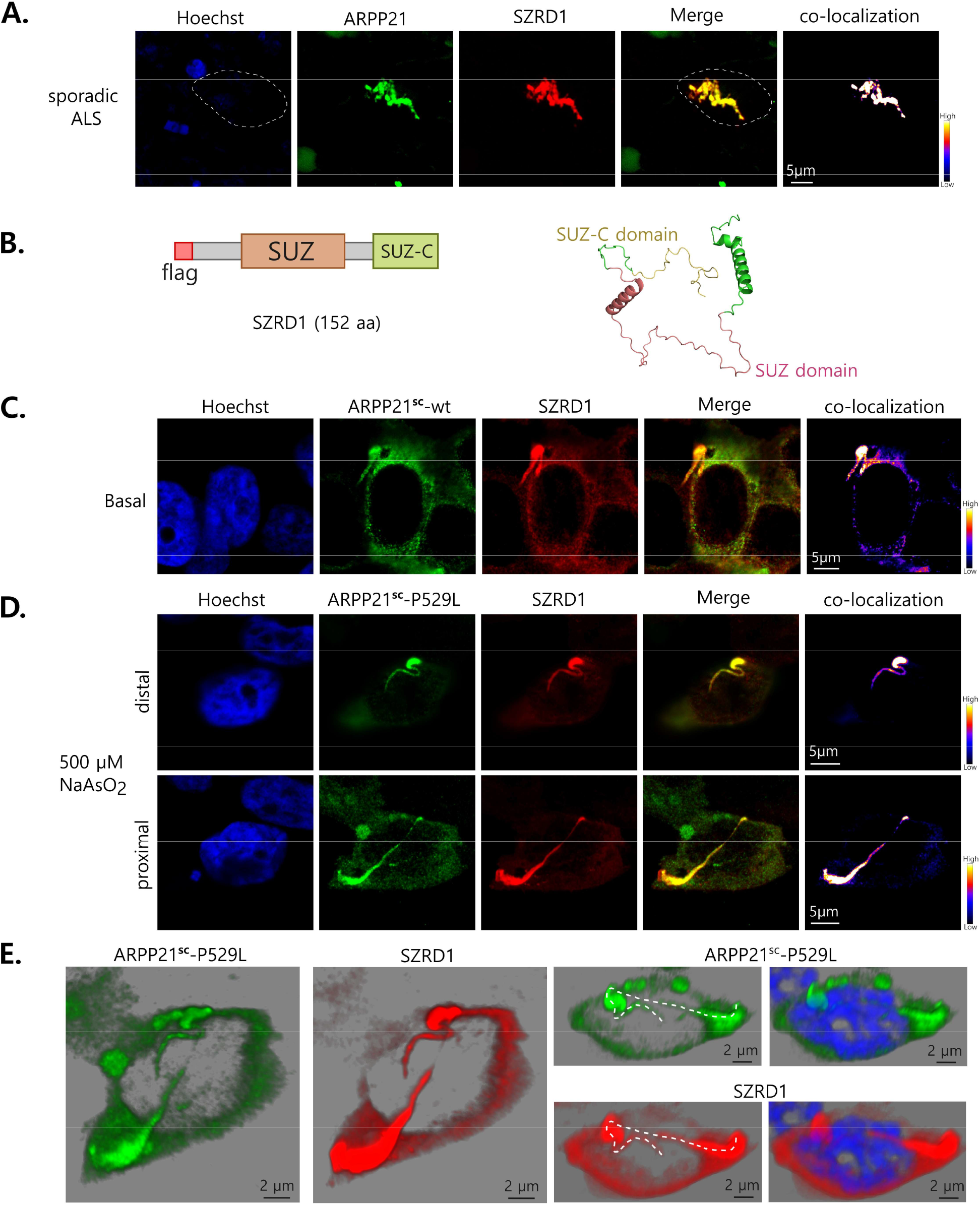
ARPP21 co-aggregates with SZRD1 in the spinal cord tissue and cultured cells. (**A**) ARPP21 aggregate co-localizes with SZRD1 in spinal cord in a case of sporadic ALS. A segmented white line demarcates the borders of aggregate-containing cell. (**B**) Schematic of flag-tagged SZRD1 (left) and Alphafold modelling of SZRD1 (right). SZRD1 is 100% disordered (MobiDB). SUZ (red) and SUZ-C (yellow) domains are highlighted in the structure. (**C, D**) ARPP21^sc^-wt and P529L variant co-aggregate with SZRD1 under basal and oxidative stress conditions in HEK293T cells, forming fibrillar structures. (**D**) A large filamentous condensate containing ARPP21^sc^-P529L and SZRD1 is observed to cross the whole cytoplasm of a cell. Distal and proximal views of the cell are shown. (**E**) 3D reconstruction of the cell and the condensate surrounding the nucleus shown on D.

AlphaFold predicted that SZRD1 has an even more disordered structure than ARPP21 (100% disordered, MobiDB; Figure 4B). When we expressed flag-tagged SZRD1 in HEK293T cells under basal conditions, we observed homogeneous expression in the cytoplasm (Supplementary Figure 5B). After stress, SZRD1 formed fibrillar structures that do not co-localize with endogenous ARPP21 (eARPP21; Supplementary Figure 5B). Co-expression of myc-tagged ARPP21^sc^ (wt, P529L or E377K variants) and flag-tagged SZRD1 in basal and oxidative stress conditions showed co-localization of the two proteins in condensates and fibrils (Figure 4C and D; Supplementary Figure 5C). Long fibrillar structures containing ARPP21^sc^-P529L and SZRD1 could even cross the whole body of the cell (Figure 4D and E). Notably, ARPP21^sc^-positive fibrils often emerged from droplet-like condensates (Figure 4C, D and E; Supplementary Figure 5C).

Together, our results show that ARPP21 is susceptible to forming spontaneous condensates and fibrils that co-localize with SZRD1, but not TDP-43, in cultured cells.

### ARPP21 has a slow dynamic in cells

To study the dynamic properties of ARPP21 condensates, we cloned and expressed clover-tagged ARPP21^sc^-wt or ARPP21^sc^-P529L and ARPP21^sc^-E377K variants, and TDP-43-wt in HEK293T cells (Figure 5A). All fluorescent chimeras formed visible condensates under basal conditions (Supplementary Figure 6A). Fluorescence recovery after photobleaching (FRAP) showed the presence of two different types of TDP-43-wt condensates that we called type I and II. Type II TDP-43-wt aggregates were the least dynamic, showing a low recovery after photobleaching (Plateau of type II TDP-43-wt = 20.56 ± 2.06%) and a slow speed of recovery (tau of type II TDP-43-wt = 9.606 ± 4.03s; Figure 5Bi). ARPP21^sc^-wt aggregates (Figure 5Bii) proved to have similar degree of recovery to type I TDP-43-wt aggregates (Plateau of ARPP21^sc^-wt = 61.99 ± 2.87%, and type I TDP-43-wt = 60.98 ± 3.44%) and higher than type II TDP-43-wt (Figure 5Bi), but a slower rate of recovery than both TDP-43 aggregates (tau of ARPP21^sc^-wt = 40.30 ± 4.89s, tau of type I TDP-43-wt = 13.25 ± 2.29s). Interestingly, even though aggregates of the ARPP21^sc^-P529L variant showed a lower recovery than ARPP21^sc^-wt (Plateau of ARPP21^sc^- P529L = 41.45 ± 5.86%; Figure 5Bii and iii), this process is not only slower compared to ARPP21^sc^-wt (tau of ARPP21^sc^-P529L = 48.29 ± 4.24s) but also to Type II TDP-43-wt aggregates, showing the lowest recovery rate constant (k) of all (k of ARPP21^sc^-P529L = 0.01756 ± 0.00396s^-1^). Aggregates formed by ARPP21^sc^-E377K variant showed a recovery similar to ARPP21^sc^-wt, but a kinetic in between ARPP21^sc^-wt and ARPP21^sc^-P529L (Plateau of ARPP21^sc^-E377K = 66.00 ± 4.13%, tau of ARPP21^sc^-E377K = 44.29 ± 7.19s; Figure 5Biv). These data support the idea that ARPP21 forms aberrant condensates that change slowly over time and that the mutant P529L has the higher pathological potential of these proteins.

We also investigated the properties of soluble ARPP21^sc^ using fluorescence correlation spectroscopy (FCS). Fluorescence intensity fluctuations of clover-tagged ARPP21 were stably fitted across multiple replicates to the autocorrelation function G(τ) using a one-component anomalous model described in the methods. This allowed the identification of an effective slow population for each ARPP21^sc^ variant (wt, P529L or E377K) and TDP-43-wt that diffuses in a mildly anomalous non-Brownian way (α<1; Figure 5C), probably due to the intracellular environment. ARPP21^sc^ variants showed a significantly lower (p<0.0001) diffusion coefficient (D for ARPP21^wt^ = 0.612 ± 0.067µm^2^/s, D for ARPP21^P529L^ = 0.625 ± 0.105µm^2^/s, and D for ARPP21^E377K^ = 0.519 ± 0.085µm^2^/s) compared to clover alone (D for clover = 58.26 ± 11.88µm^2^/s), and similar to cytoplasmic TDP-43-wt (D for cytoTDP-43-wt = 0.520 ± 0.043µm^2^/s), indicating very slow diffusion related to oligomerization, complex formation, or condensate-like behavior. Molecular brightness (MB), a parameter that is stoichiometry-sensitive, suggests that, on average, a transient mixture of small oligomers for ARPP21^sc^ (wt and variants) and a more assembled/multimerized form for TDP-43-wt exists in the cytoplasm (ARPP21^sc^-wt = 1.715 ± 0.393, ARPP21^sc^-P529L = 1.593 ± 0.379 and ARPP21^sc^-E377K = 1.770 ± 0.169; TDP- 43-wt = 2.36 ± 0.274; Figure 5C). Together, FCS results support a model of a heterogenous population of soluble ARPP21 that behaves like a continuum of slow mobility states with transient interactions.

Additionally, we performed an analysis of the droplet state of the proteins using FuzDrop (*29–31*) (https://fuzdrop.bio.unipd.it/predictor). We obtained a residue-based droplet-promoting probabilities (P_DP_) graph for TDP-43-wt, ARPP21^sc^-wt, and the variants, and the probability of spontaneous liquid-liquid phase separation (P_LLPS_) for those proteins. The values of P_LLPS_ for TDP-43-wt, ARPP21^sc^-wt, ARPP21^sc^-P529L, and ARPP21^sc^-E377K were 0.8981, 0.9992, 0.9994, and 0.9994, respectively (Supplementary Figure 6B). This analysis supported our experimental evidence of biophysical differences between TDP-43-wt and the ARPP21^sc^ variants.

In summary, these experiments suggest that ARPP21 forms aberrant condensates, which are preceded by a spectrum of slow mobility and dynamics. These results suggest that ARPP21^sc^ spend relatively less time in mobile oligomeric intermediates, but once nucleated, rapidly converts into stable low-exchange assemblies.

### ARPP21 induces toxicity in motor neurons

Next, we studied whether ARPP21^sc^-wt and ARPP21^sc^-P529L variants induce a toxic effect in a cell model relevant to ALS. When we transduced human iPSC-derived motor neurons (MNs) with AAV9 carrying myc-tagged versions of ARPP21^sc^-wt or ARPP21^sc^-P529L, we observed cytoplasmic condensates under basal conditions, unlike with AAV9-GFP. These ARPP21 condensates rarely co-localized with endogenous TDP-43 (Figure 6A). Under oxidative stress and without virus transduction, we noticed the formation of endogenous ARPP21 (eARPP21)- positive condensates, which largely did not co-localize with TDP-43 (Figure 6B). Then, we transfected human cultured neuronal cells (SH-SY5Y) with myc- ARPP21^sc^ constructs and performed ATP viability assays. We observed that ARPP21^sc^ overexpression significantly reduced the survival of cells, an effect that was slightly stronger for P529L and E377K variants (Figure 6C). The transduction of AAV9- ARPP21^sc^-wt or - ARPP21^sc^-P529L in human iPSC-derived motor neurons also reduced the viability of the cells compared to the AAV9-GFP transduced control (Figure 6D).

**Figure 5.**
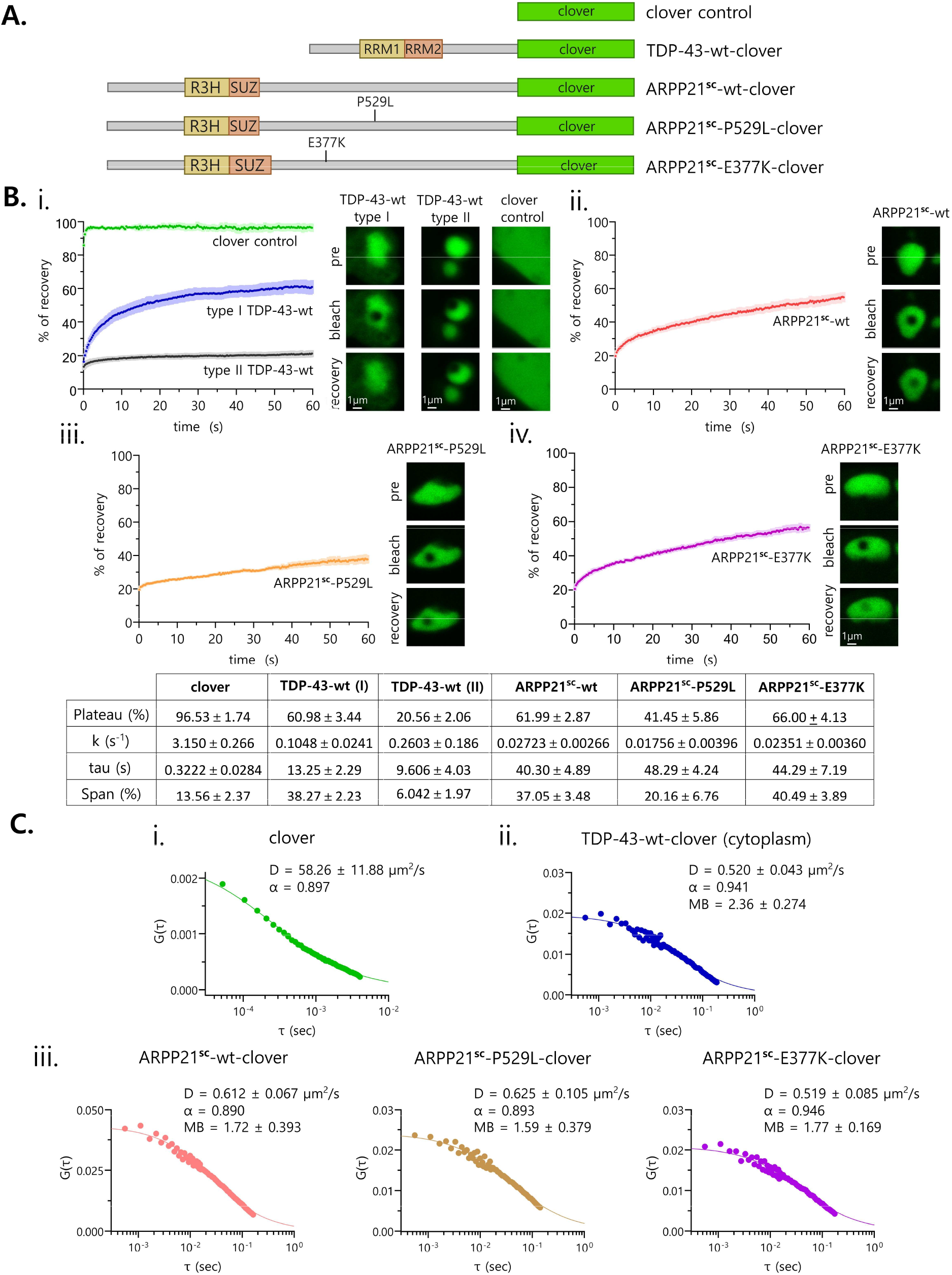
ARPP21 has a slow dynamic. (**A**) Schematic of the clover-tagged constructs used for fluorescence recovery after photobleaching (FRAP) and fluorescence correlation spectroscopy (FCS) experiments in HEK293T cells. (**B, i**) Two types of TDP-43-wt condensates were observed by FRAP. TDP-43- wt type I condensates are more dynamic than type II. (**A, ii**) Fluorescence of ARPP21^sc^-wt condensates recover at a slower rate than TDP-43 condensates. (**A, iii**) Condensates of ARPP21^sc^-P529L variant recovers at a slower rate than ARPP21^sc^-wt’s and are the least dynamic of all condensates (**A, iv**) ARPP21^sc^-E377K variant forms condensates that are slightly less dynamic than ARPP21^sc^-wt condensates. FRAP parameters for the different aggregates are shown in the bottom table. (**C**) FCS autocorrelation curves for ARPP21^sc^-wt, and variants P529L and E377K, compared to clover and TDP-43-wt. ARPP21^sc^ variants have a similar diffusion (diffusion coefficient, D) to TDP-43-wt and two orders of magnitude slower than clover alone. All the proteins show a similar anomalous component (α), probably due to a similar cellular environment. Molecular brightness (MB) values of ARPP21^sc^ variants support the formation of smaller oligomers/complexes than TDP-43.

**Figure 6.**
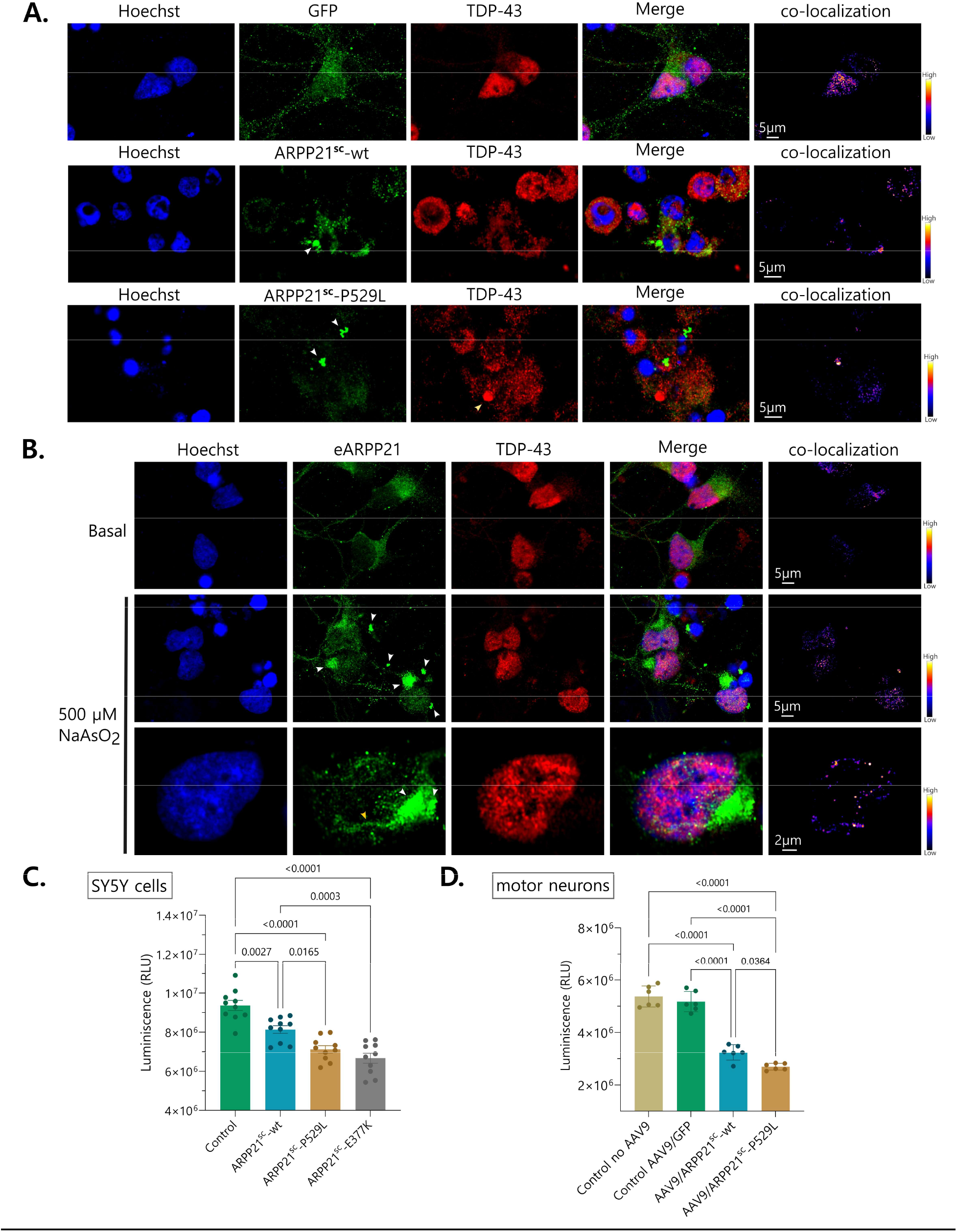
ARPP21 induces cell toxicity and death in motor neurons. (**A**) Human iPSC-derived motor neurons were transduced with AAV-9/GFP (control), AAV9- ARPP21^sc^-wt, or AAV9/ ARPP21^sc^-P529L and maintained under basal conditions. GFP shows a fine granular and homogenous distribution (top panel). ARPP21^sc^-wt (middle panel) and P529L (bottom panel) both form condensates in the cytoplasm (white arrowheads) that rarely co- localize with endogenous TDP-43. Condensates of TDP-43 without ARPP21 are also observed (yellow arrowhead). (**B**) Oxidative stress also induces the formation of condensates of endogenous ARPP21 (eARPP21) (white arrowheads) and fibrils of ARPP21 sometimes observed emerging from condensates (third panel) (yellow arrowhead). Rare and low-level co-localization with TDP-43 is observed in ARPP21 condensates. (**C**) Cell viability was reduced in SH-SY5Y neuronal cells transfected with vectors expressing ARPP21^sc^-wt, ARPP21^sc^-P529L and ARPP21^sc^-E377K. Empty vector was used as a control for transfection. (**D**) Cell viability was also reduced in human iPSC-derived motor neurons transduced with AAV9/ARPP21^sc^-wt or AAV9/ARPP21^sc^-P529L. Transduction with AAV-9/GFP was used as control. ARPP21^sc^ variants show a stronger effect on diminishing survival than ARPP21^sc^-wt.

Together, our results indicate that ARPP21 forms spontaneous fibrils and condensates in cultured motor neurons under basal conditions and reduces the viability of neuronal cells and motor neurons.

### ARPP21 spreads by inducing TNT formation

In cultured cells and spinal cord tissue, we observed the formation of ARPP21-positive bridge-like structures of different lengths resembling tunneling nanotubes (TNTs), membrane protrusions that allow cell-cell long-range communication through the transfer of molecules and organelles (*32*). TNT presence was confirmed by immunostaining of HEK293T cells with F-actin antibodies, a common marker of these structures (Figure 7A). We observed that F-actin-positive TNTs connect the cytoplasm of two cells and transport condensates of ARPP21^sc^-wt and ARPP21^sc^-P529L (Figure 7A, B, C and D; Supplementary Figure 7A, B). The free-floating hallmark of the TNTs was evident after Z-stack imaging and 3D orthogonal reconstruction (Figure 7B). Co-immunostaining showed that SZRD1 is also transported through TNTs and co- localizes with ARPP21^sc^ (Figure 7A and B; Supplementary Figure 7B and C). Often, we observed ARPP21^sc^ condensates enlarged and accumulated at the terminal docking region of the TNT (Figure 7C and Supplementary Figure 7D). TDP-43 instead localized in TNTs to a lesser extent and mainly after stress, but TDP-43-positive granules were observed separated from ARPP21^sc^ (Figure 7C). Only in the cytoplasm of the recipient cell were stress-induced TDP-43 condensates observed in proximity to ARPP21^sc^ condensates but without colocalization (Figure 7D). In the dense tissue of the human spinal cord, we frequently observed F-actin-positive TNT-like structures (*33, 34*) that contained ARPP21, sometimes with a reminiscent TNT docking region (Supplementary Figure 7E and F).

**Figure 7.**
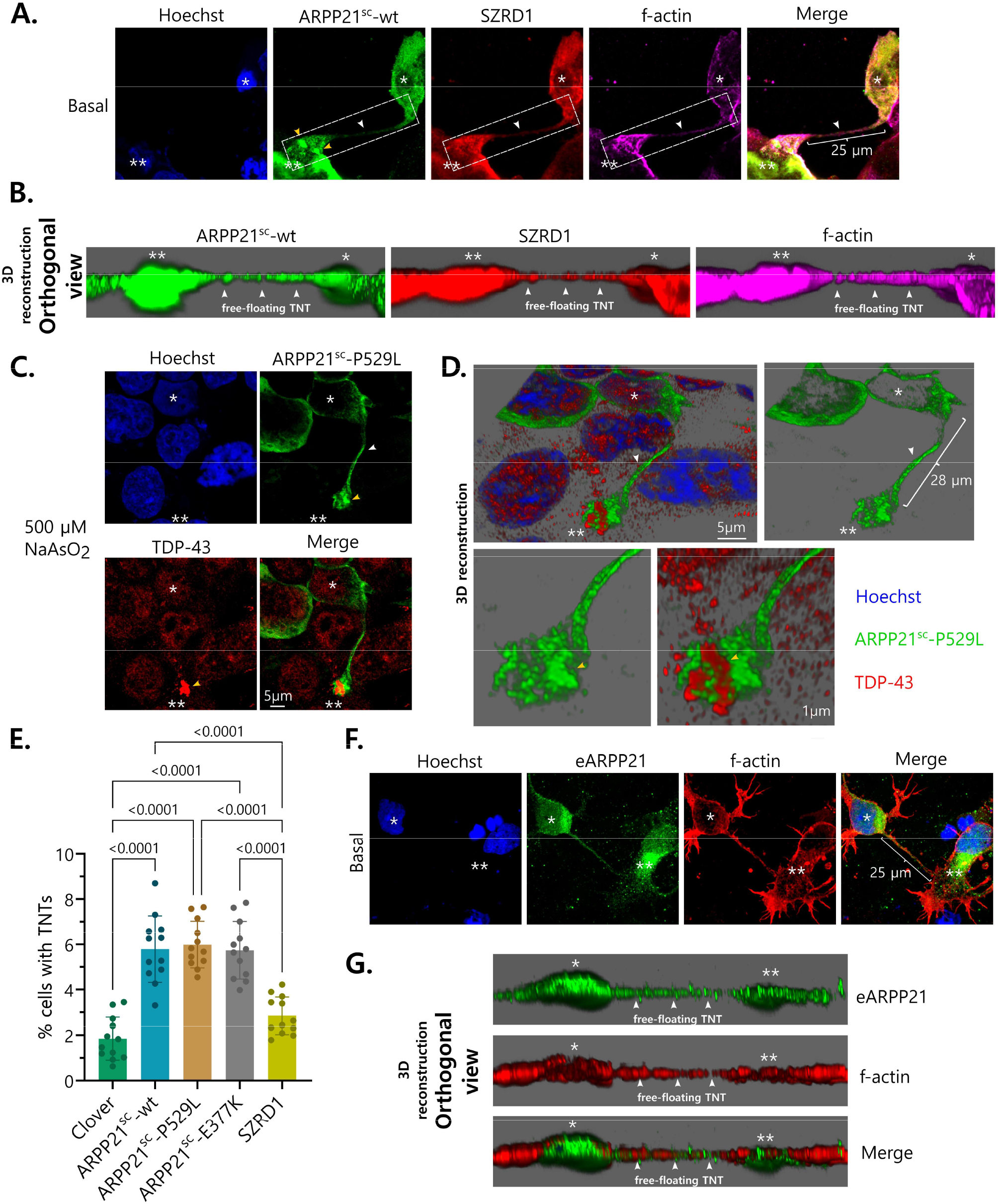
ARPP21 spreads through tunnelling nanotubes (TNTs) between cells. (**A**) F-actin-positive TNT contains myc-tagged ARPP21^sc^-wt and flag-tagged SZRD1 in HEK293T cells under basal conditions. (**B**) orthogonal view of 3D reconstruction showing free-floating F-actin-positive TNT (white arrowheads) with ARPP21^sc^ and SZRD1 condensates from the cell shown in A. (**C**) TNT (white arrowhead) that transports ARPP21^sc^-P529L shows the formation of larger ARPP21^sc^ and TDP-43 condensates without co-localization in the recipient cell under oxidative stress (yellow arrowheads). (**D**) 3D reconstruction from cell in C shows that ARPP21^sc^-P529L condensates in the TNT (white arrowhead) are close but do not merge with TDP-43 condensates (yellow arrowheads) in the recipient cell. (**E**) Quantification of TNTs shows an increase in TNT-containing cells after ARPP21^sc^ transfection (wt and variants) compared to clover and SZRD1 controls. (**F**) Human iPSC-derived motor neurons also show F-actin-positive TNTs containing endogenous ARPP21 (eARPP21). (**G**) 3D orthogonal view of the cell on F shows the presence of free-floating TNT. Donor (one asterisk) and recipient cells (two asterisks) are highlighted in A, B, C, D, F and G.

The observation of TNTs in HEK293T cells transfected with ARPP21^sc^ prompted us to investigate whether ARPP21 could induce TNT formation. Cells transfected with ARPP21^sc^ showed an increase in the percentage of TNT-connected cells compared to clover and SZRD1 controls; no difference was observed between ARPP21^sc^ -wt, -P529L, and -E377K variants (Figure 7E). F-actin-positive free-floating TNTs containing eARPP21 were also occasionally observed in iPSC-derived MNs under basal conditions (Figure 7F and G).

Our data suggest that ARPP21^sc^ expression induces the formation of TNTs to facilitate the transport and spreading of the protein between cells.

### Rare *ARPP21* variants exist in ALS

Upon initial curation by the ClinGen ALS Gene Curation Expert Panel (GCEP), the evidence regarding the association between the gene *ARPP21* and ALS was considered “limited”, which is a classification given to genes that have been proposed to be associated with or causative of ALS in the literature but still have insufficient data to definitely prove the relationship. However, a recent large-scale ALS case-control analysis (individuals with ALS = 13,138, controls = 69,775) investigated two specific variants within *ARPP21*, namely p.P563L and p.P747L, and identified strong, significant associations with ALS, with both variants displaying odds ratios > 40 (*13*). These results implicate the gene as an important risk factor for ALS.

To compliment these findings and investigate other variants that may be of relevance for ALS in *ARPP21*, we extracted rare variant carrier counts from the individuals with ALS and controls in the ALS Knowledge Portal and Project MinE ALS Consortium datasets (Table 1). Interestingly, when the variants identified within the two datasets were compared to those previously curated as part of the ClinGen ALS GCEP assessment of *ARPP21*, no variants in common were identified (Figure 8; Supplementary Figure 8). We also reviewed the dataset for the two variants recently proposed as associated with ALS in the large-scale case-control analysis and found that while p.P563L was not identified in the ALS Knowledge Portal nor Project MinE ALS Consortium datasets, p.P747L was carried by eight individuals with ALS and no controls across the two datasets. Of note, p.P563L is known to have sub-par call rates in some exome cohorts.

**Figure 8.**
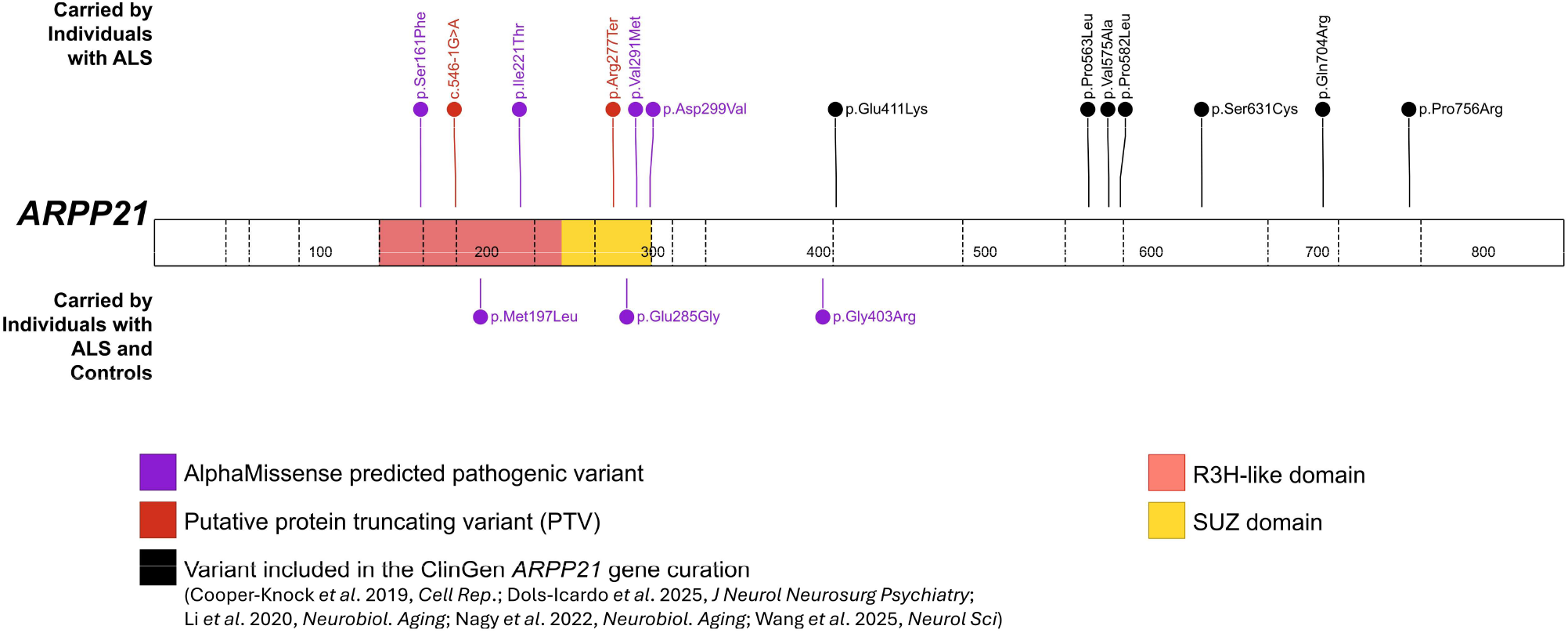
Rare, non-synonymous variants of potential interest were identified in *ARPP21*. Analyses were performed with the ALS Knowledge Portal (ALSKP; individuals with ALS = 3864, controls = 7839) and Project MinE ALS sequencing consortium (individuals with ALS = 6596, controls = 2454) datasets. Variants only identified in controls were excluded from the figure (n=39). Seven missense variants from four peer-reviewed manuscripts are also indicated that were curated by the ClinGen Gene Curation Expert Panel (GCEP) in the review of *ARPP21*, none of which were predicted to be pathogenic by AlphaMissense.

**Table 1.** Rare variant counts identified in *ARPP21* in the ALS Knowledge Portal (ALSKP; individuals with ALS = 3864, controls = 7839) and Project MinE ALS sequencing consortium (ProjMinE; individuals with ALS = 6596, controls = 2454) datasets.

| Variant Type | ALS –<br>ALSKP (n) | Controls –<br>ALSKP (n) | ALS –<br>ProjMinE (n) | Controls –<br>ProjMinE (n) |
| --- | --- | --- | --- | --- |
| Synonymous | 7 | 22 | 3903* | 1447* |
| Missense | 98 | 210 | 165 | 48 |
| AlphaMissense predicted pathogenic | 0 | 2 | 7 | 8 |
| Putative PTVs | 0 | 1 | 2 | 1 |

\*Largely driven by a single synonymous variant. Abbreviations: PTVs, protein truncating variants.

## Discussion

While a small set of recurrent pathological hallmarks of ALS is widely accepted, the disease remains mechanistically heterogeneous and incompletely understood. The emergence of *ARPP21* which not only exhibits deleterious variants in ALS (*10–13*) but also shows a broad pathological distribution, demonstrates how limited our knowledge is about the complex pathophysiology of this neurodegenerative disease.

Historically, SOD1 has been considered excluded from TDP-43 pathology in ALS. However, several studies have shown that both pathologies can coexist in the same patient and co-deposit in the same inclusions (*35–37*). Our study reveals a picture in which not every protein that frequently forms aggregates in ALS (familial and sporadic) is connected to TDP-43 pathology or shares condensate similarities, even if it is an RBP. ARPP21 fibrillar aggregates show a singular morphology that to us resembles “growing crystals” and in early stages in cells behave like dense compartments. It is noteworthy that together with the segregated nature of ARPP21 and TDP-43 aggregates (Figure 1), ARPP21 aggregates in motor neurons without TDP-43 pathology (Figure 2), including cells that would previously have been considered “healthy” in the ALS tissue. This suggests that ARPP21 might be an earlier marker of motor neuron disease, an idea that warrants further research.

ARPP21 was more prone to aggregate formation than TDP-43, showing condensates that are less dynamic in cells. This suggests that ARPP21 might bypass prolonged dynamic oligomeric states, transitioning efficiently into kinetically trapped states, whereas TDP-43 could spend more time in reversible/mobile oligomeric states. This critical difference could be the molecular basis of the pathological segregation between ARPP21 and TDP-43 in ALS.

It is important to note that aggregation of proteins such as ARPP21 and TDP-43 is governed in cells by a set of overlapping conditions. Sequence determinants (intrinsic nucleation potential), stability and folding kinetics (conformation accessibility), phase behavior and supersaturation (thermodynamic driving force), and cellular proteostasis capacity, all together play crucial roles in the process (*38–40*). IDRs and aggregation-prone regions (APRs), short sequences with β- aggregation propensity, are part of the intrinsic aggregation potential. The exposed disordered area of a protein, which is in contact with other proteins and RNAs in the cell, could also be included here. ARPP21 has a higher percentage of disorder and a higher tendency to form condensates than TDP-43 (% disorder ARPP21= 87.3, TDP-43= 60.6; probability of spontaneous liquid-liquid phase separation (P_LLPS_) for ARPP21 = 0.9992 and TDP-43 = 0.8981) and also doubles the size of TDP-43 (scARPP21= 813aa; TDP-43 = 414 aa). All of this, together with the localization of ARPP21 that is mainly cytoplasmatic, where the RNA concentration is too low to properly regulate phase separation (*41*), could make ARPP21 more sensitive to intracellular changes and explain at least in part its aggregation-prone behavior in more rigid condensates.

The non-ubiquitinated nature of ARPP21 aggregates is also intriguing. Intracellular aggregates in neurodegeneration show at the very least different degrees of ubiquitination at some stage of the disease. For example, mtSOD1 aggregates are partly oligo-ubiquitinated (*42*) and huntingtin polyQ aggregates precede ubiquitination in ALS and Huntington disease mouse models, respectively (*43*). While ubiquitin-positive aggregates have been considered a hallmark in ALS motor neurons (*44*), our results suggest that ubiquitination might be less extensive among neuropathological-associated aggregates than was previously thought. The fact that increased levels of ARPP21 induce condensate formation and cell toxicity suggests that other mechanisms different than the ubiquitin/proteasome, such as disaggregation and heat shock chaperons, could be involved in the degradation of these non-ubiquitinated aggregates (*45–48*).

The spreading of ARPP21 condensates between cells through TNTs constitutes a highly efficient way of reducing cellular stress by distributing the overload of highly disordered proteins and relieving the burden of aggregate accumulation. This might be the case for α-synuclein, which is transferred between neuronal and microglial cells using TNTs, possibly as a rescue mechanism (*49*). TNTs and other intercellular bridges are essential for the physiology of the cell because their ability to transport biological cargo serves to share not only proteins but also RNAs, ions and organelles (*32, 49, 50*). It could be tempting to consider TNTs as potential therapeutic targets to stop the spread of proteins such as ARPP21. However, due to their critical physiological relevance, TNTs are structures that are very challenging to manipulate directly without affecting healthy cells. This idea favors intervening upstream at the aggregation step, which has the added advantage of limiting the aggregate-induced nanotube formation that drives propagation in the first place.

In summary, we have described that ARPP21, a SUZ domain-containing RNA-binding protein, defines an uncovered parallel pathological axis independent of TDP-43 pathology. We found that ARPP21 forms low dynamic condensates more efficiently than TDP-43 and spreads through cells, inducing the formation of TNTs. We described that the P529L spinal cord variant (P653L in the full length sequence of ARPP21) is more pathological than the wt isoform, which enhances its clinical relevance. Our genomic analysis supports the need for broader investigations of *ARPP21* variants in individuals with ALS.

## Acknowledgements

This work was supported by a generous donation from the Temerty Family Foundation. M.J.S. is supported by the Canadian Institutes of Health Research (CIHR). We sincerely thank the Mendoza family from Chile who reached out to us to better understand the ALS-associated mutation present in their family. Their commitment to advancing knowledge of this condition was a major source of inspiration for this work.

## Author contributions

D.C.M and C.A.D. conceived the project, performed and analyzed the experiments, performed the theoretical analyses, analyzed and checked the overall quality of the data, prepared the illustrations, and drafted the manuscript; A.A.D and S.M.K.F performed the genetics analysis and prepared the genetics-related illustrations and text; M.J.S. secured funding, reviewed and provided valuable input for the final version of the manuscript. All authors reviewed and approved the final version of the manuscript.

## Generative AI statement

The authors declare that generative AI was not used in the creation of this manuscript.

---

**Supplementary Figure 1.**
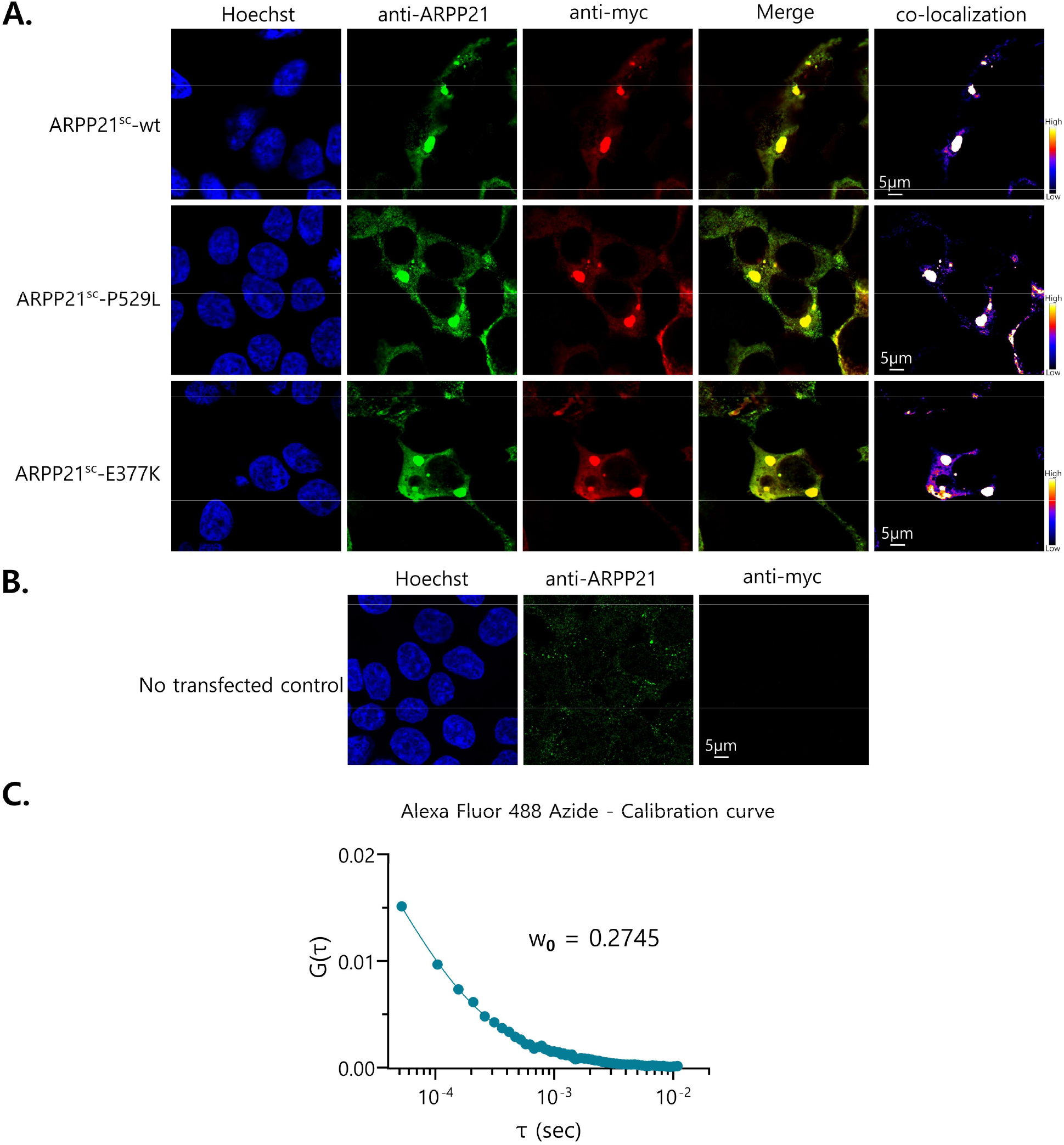
ARPP21 antibody characterization and FCS calibration curve. (**A**) Immunofluorescence using rabbit anti-ARPP21 and mouse anti-myc on HEK293T cells transfected with plasmid expressing myc-tagged ARPP21^sc^-wt, ARPP21^sc^-P529L, and scARPP21^sc^-E377K. Complete co-localization between the two channels indicates that both antibodies recognize the same protein. (**B**) Negative control of immunofluorescence using anti- ARPP21 and anti-myc on non-transfected HEK293T cells. The anti-myc antibody is not able to recognize the endogenous ARPP21 expressed in low levels in the cells. (**C**) FCS calibration curve for Alexa 488 Azide used to determine the w_0_ value of our instrument.

**Supplementary Figure 2.**
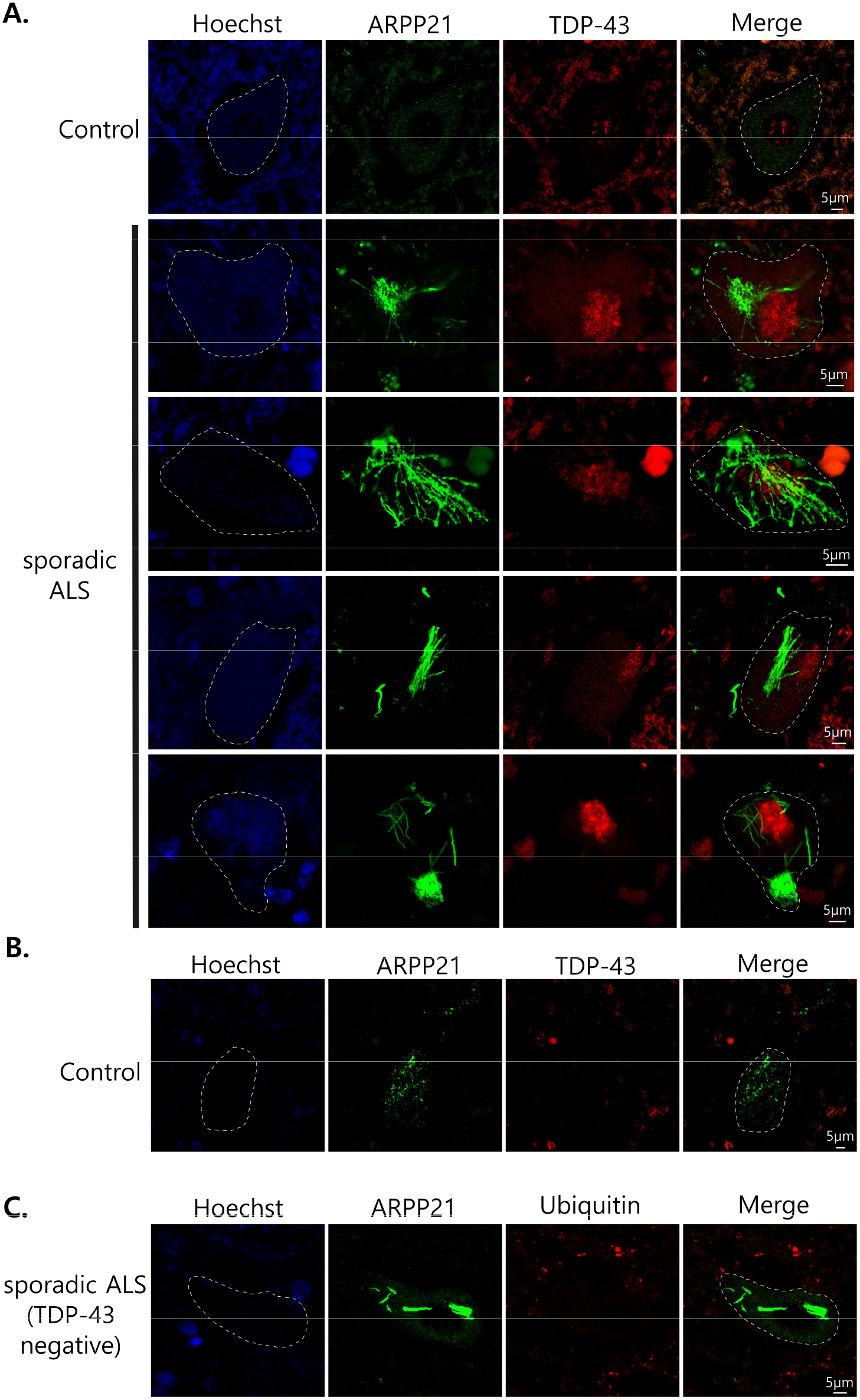
ARPP21 forms aggregates in spinal cord motor neurons without TDP-43 pathology in sporadic ALS. (**A**) ARPP21 forms fibrillar aggregates that do not co-localize with TDP-43 and are observed in neurons without TDP-43 pathology. Control cells (upper panel) do not show ARPP21 aggregates. (**B**) Occasionally, larger granules of ARPP21 are observed in controls without fibril formation. (**C**) ARPP21 fibrils do not co-localize with ubiquitin in a case negative for TDP-43 pathology. A segmented white line demarcates the borders of neurons.

**Supplementary Figure 3.**
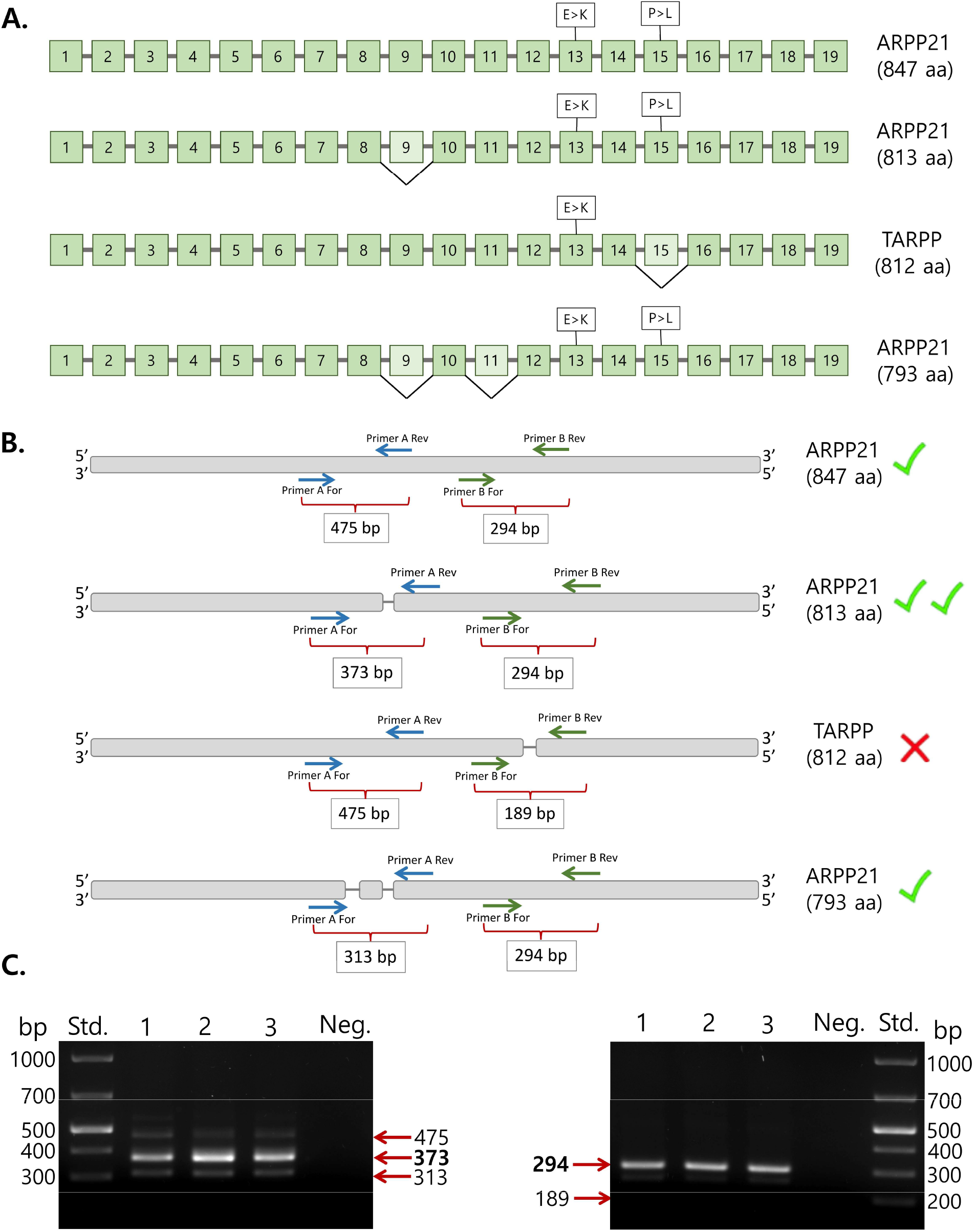
Schematic of ARPP21 differentially spliced transcripts. (**A**) Four ARPP21 isoforms were previously reported in human tissues. ARPP21^847aa^ is the full-length version of the protein; ARPP21^813aa^ isoform lacks exon 9; ARPP21^812aa^ (TARPP) lacks exon 15; and ARPP21^793aa^ lacks exons 9 and 11. (**B**) Expected amplicons for two sets of primers (A and B) used to detect the different ARPP21 isoforms in human spinal cord by PCR. (**C**). Agarose gels with PCR products of the set of primers A (left panel) and set of primers B (right panel) showing that ARPP21^847aa^, ARPP21^813aa^ and ARPP21^793aa^ are all expressed in spinal cord, but ARPP21^813aa^ is the most abundant splicing variant (373 bp amplicon with set of primers A). ARPP21^812aa^ (TARPP) was not detected (band at 189 bp with the set of primers B is absent).

**Supplementary Figure 4.**
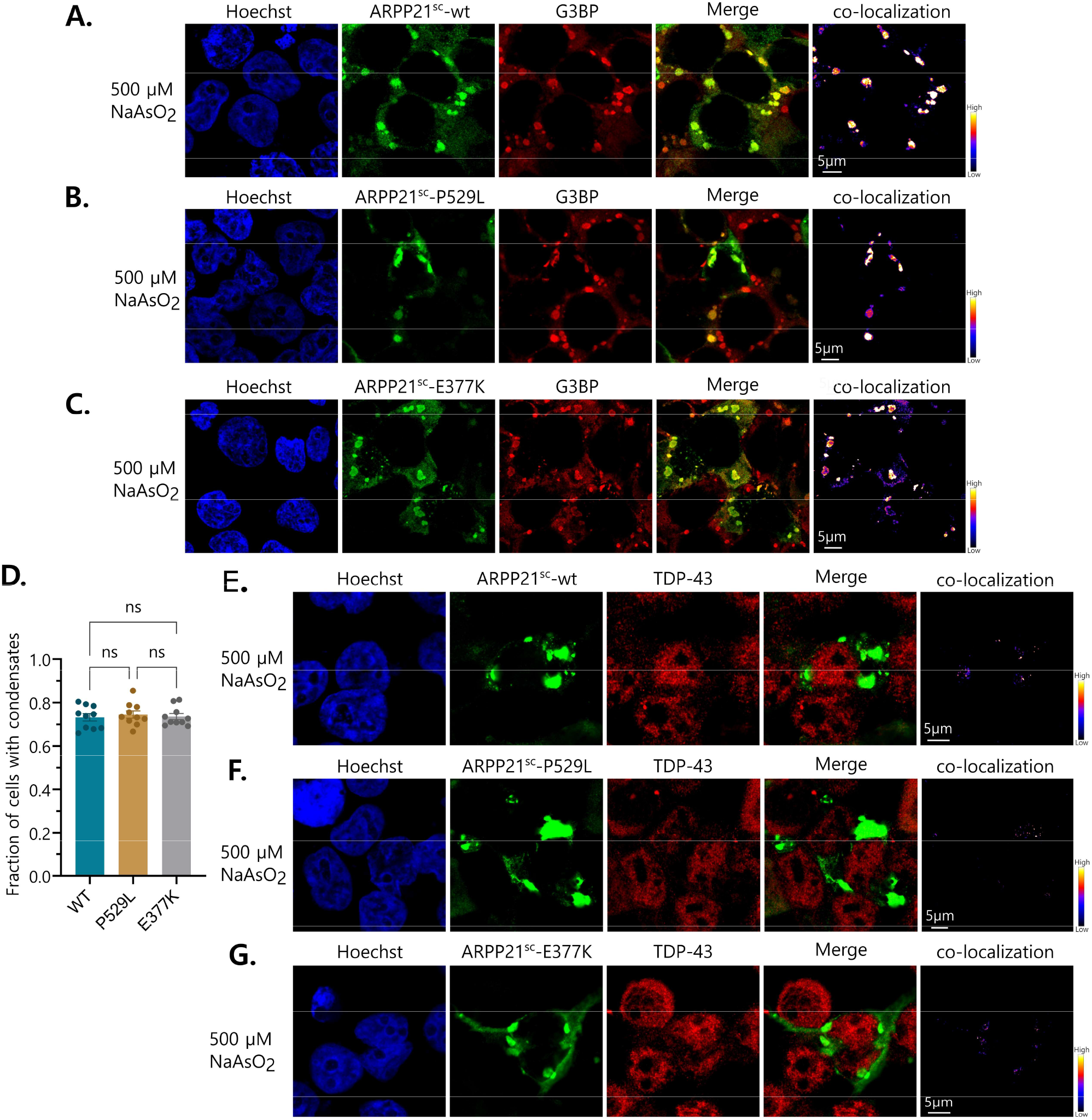
ARPP21 aggregation in cells after stress. (**A, B** and **C**) ARPP21^sc^-wt, -P529L and -E377K condensates co-localize with G3BP under oxidative stress induced by NaAsO_2_ in HEK293T cells. (**D**) Quantification of condensates in A, B and C. (**E**, **F, G**) Condensates of ARPP21^sc^-wt, -P529L and -E377K rarely co-localize with TDP-43 after stress.

**Supplementary Figure 5.**
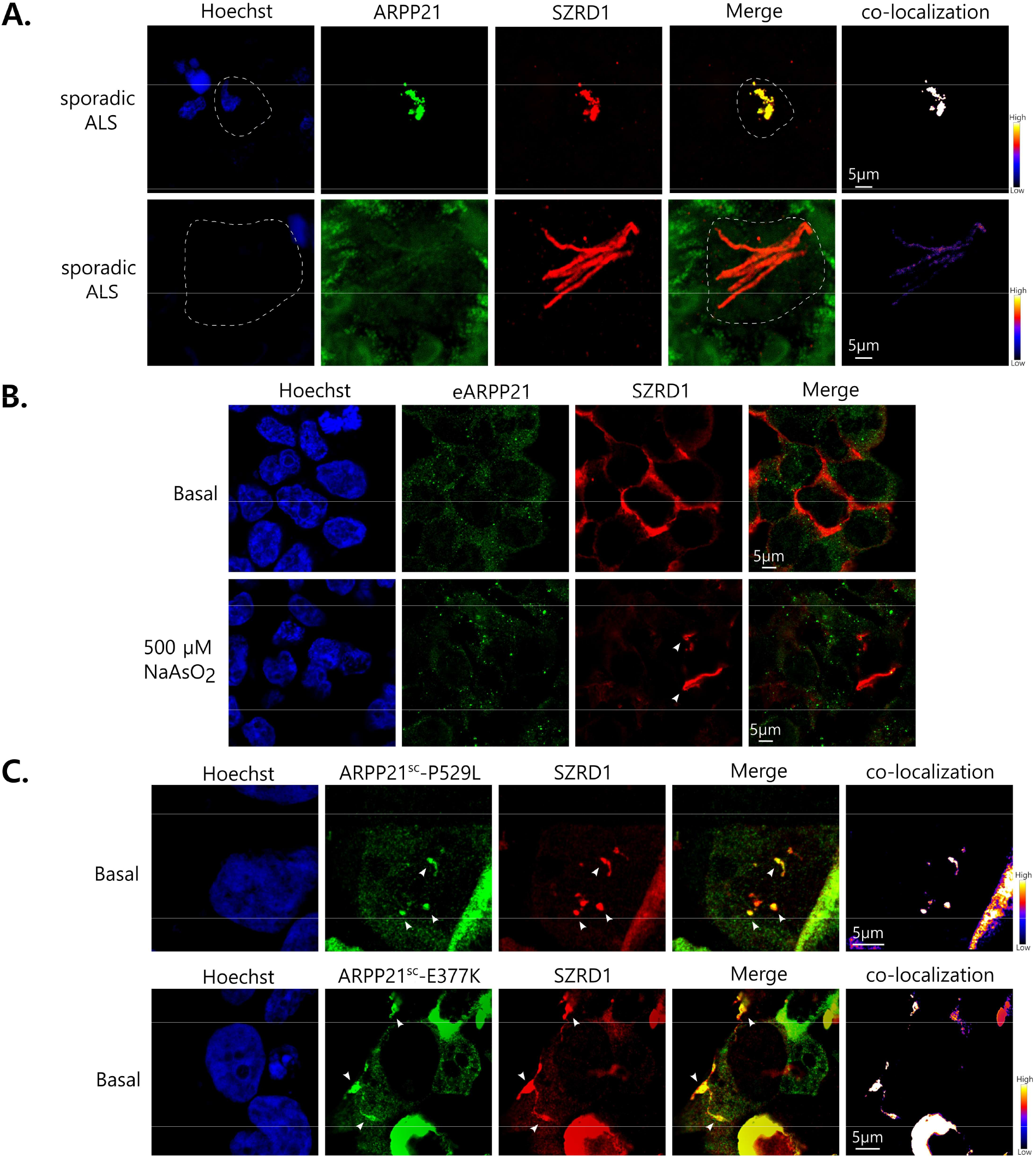
ARPP21 co-localizes with SZRD1 in spinal cord tissue and cells. (**A**) ARPP21 co-localizes with SZRD1 in sporadic ALS spinal cord, forming amorphous aggregates or fibrils. A segmented white line demarcates the borders of aggregate-containing cells. (**B**) Endogenous ARPP21 (eARPP21) granules rarely co-localize with transfected SZRD1 in HEK293T cells under basal and stress conditions; even when under stress, SZRD1 forms fibrils. (**C**) Spontaneous ARPP21^sc^-P529L and ARPP21^sc^-E377K globular and fibrillar condensates co-localize with SZRD1 under basal conditions.

**Supplementary Figure 6.**
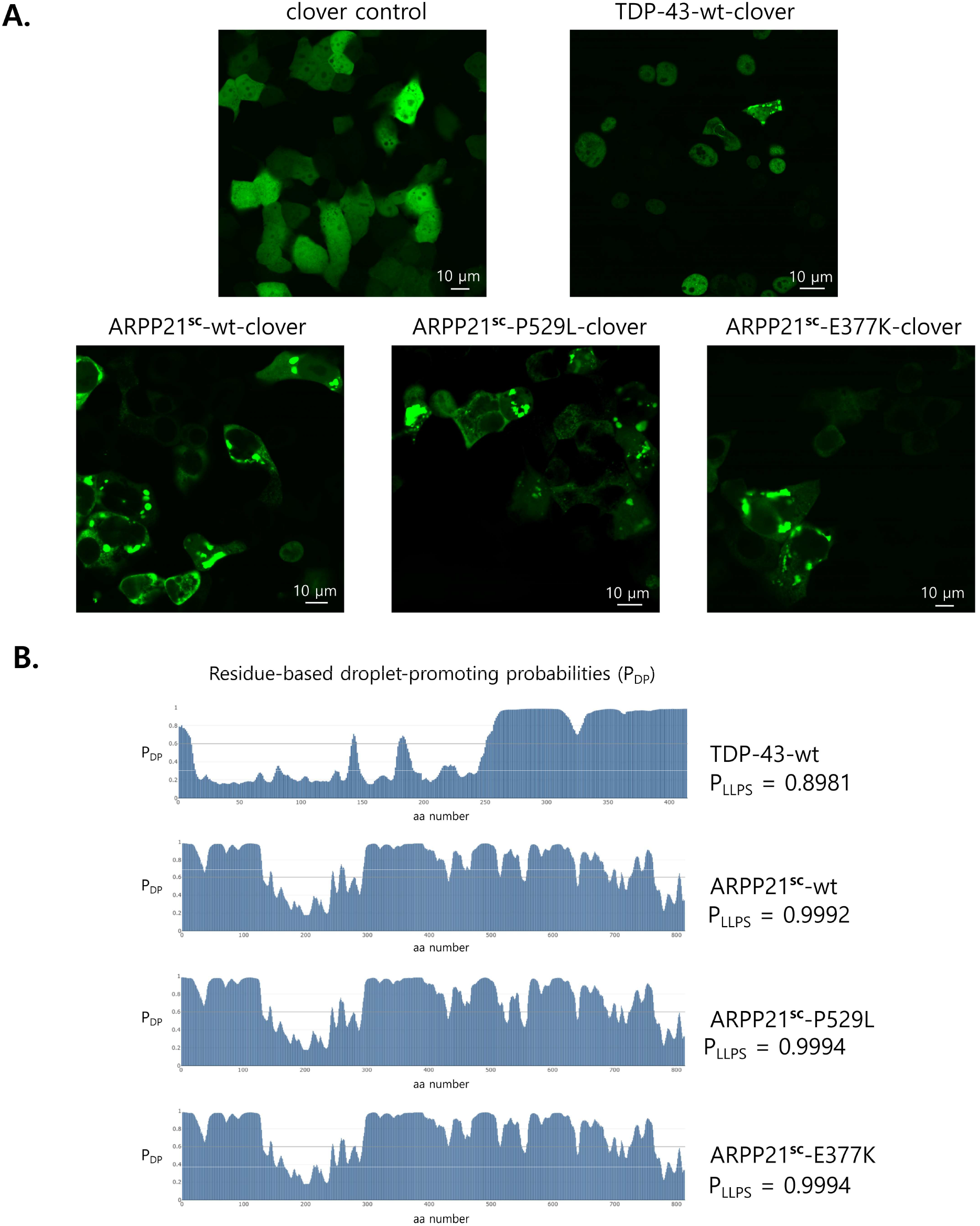
Clover-tagged ARPP21 constructs form spontaneous condensates. (**A**) Clover-tagged ARPP21^sc^-wt, ARPP21^sc^-P529L, ARPP21^sc^-E377K, and TDP-43-wt form spontaneous condensates in the cytoplasm of HEK293T cells that were used to analyze biophysical properties using FRAP. The FCS analysis was performed in regions of the cells that did not show condensate formation. (**B**) Analysis of the droplet state of the proteins using FuzDrop. The residue-based droplet-promoting probabilities (P_DP_) graphs for TDP-43-wt, ARPP21^sc^-wt, ARPP21^sc^-P529L, and ARPP21^sc^-E377K are shown. Also, it shows the probability of spontaneous liquid-liquid phase separation (P_LLPS_) for the proteins.

**Supplementary Figure 7.**
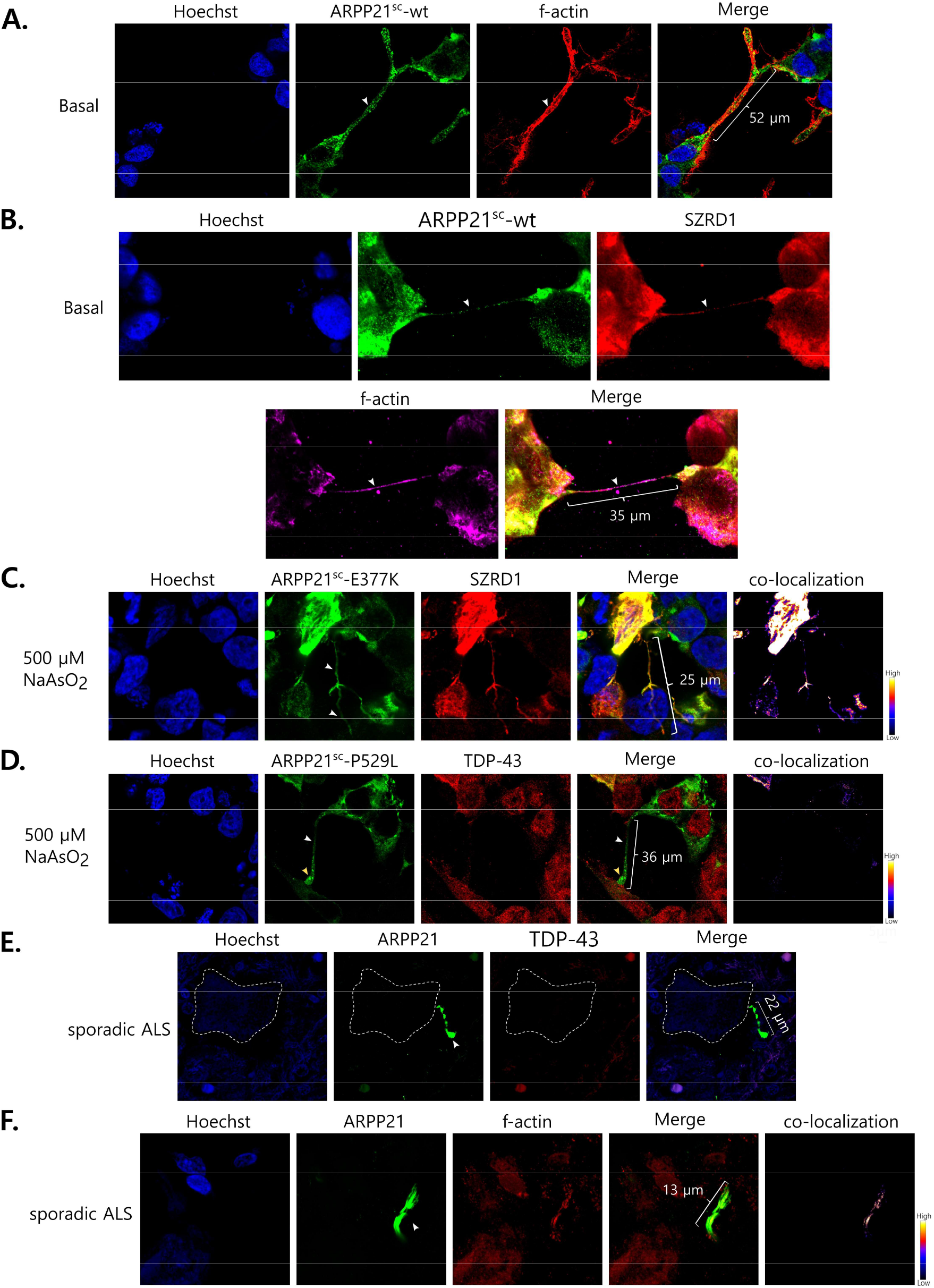
ARPP21 and SZRD1 are transported between cells through TNTs. (**A**) F-actin-positive TNT containing ARPP21^sc^-wt under basal conditions in HEK293T cells. (**B**) F-actin-positive TNT containing ARPP21^sc^-wt and SZDR1 under basal conditions. (**C**) TNT (white arrowheads) containing ARPP21^sc^-E377K and SZRD1 under oxidative stress. (**D**) TNT (white arrowhead) containing ARPP21^sc^-P529L under oxidative stress shows large condensates close to the recipient cell (yellow arrowhead). TDP-43 was rarely observed in TNTs. (**E**) ARPP21-positive TNT-like structures in human spinal cord tissue of sporadic ALS cases do not co-localize with TDP-43. A structure reminiscent of the terminal docking region of a TNT is observed (yellow arrowhead). A segmented white line demarcates the borders of the motor neuron. (**F**) ARPP21-positive TNT-like structures in human spinal cord tissue of sporadic ALS co-localize with F-actin. TNT-like structures are indicated with white arrowheads.

**Supplementary Figure 8.**
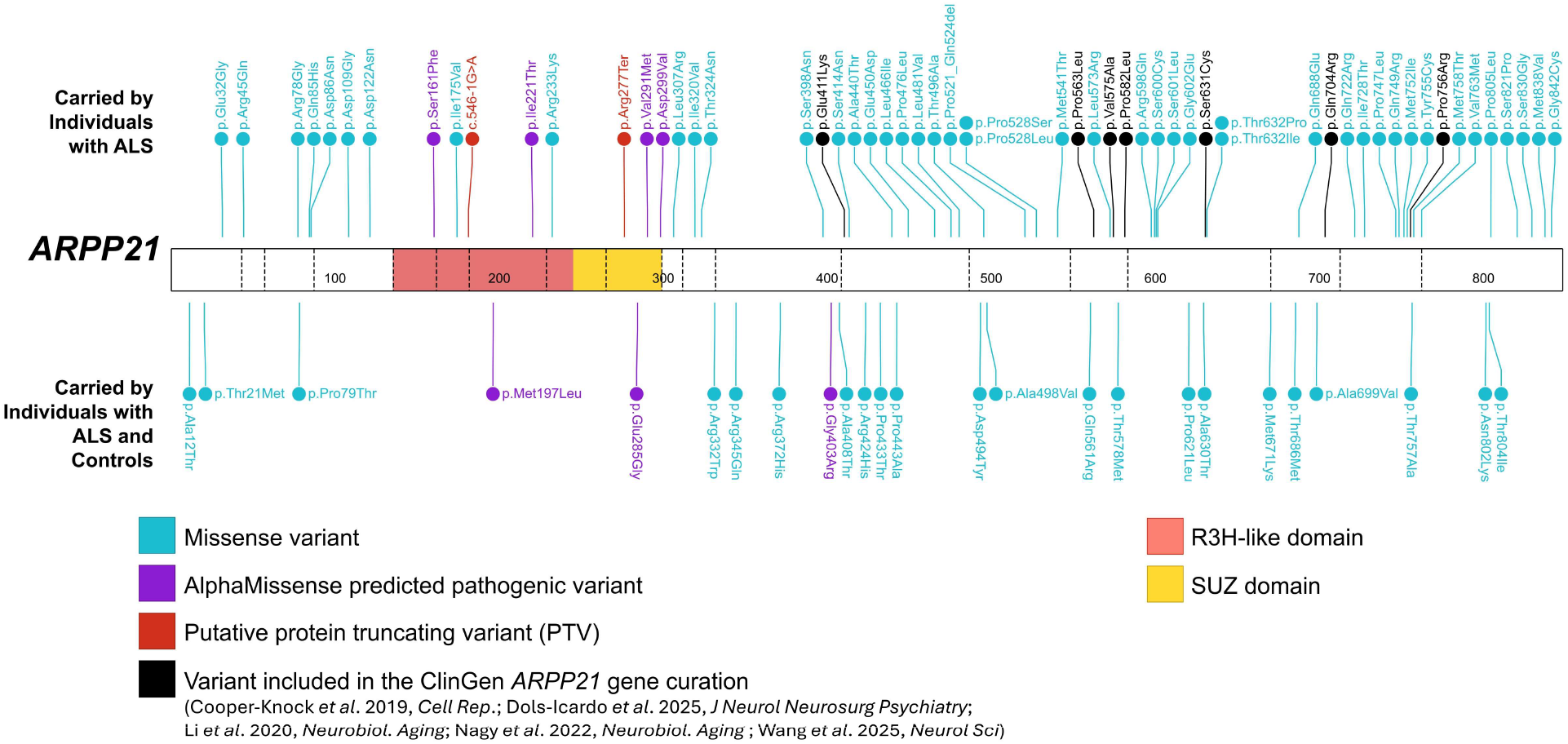
**All rare, non-synonymous variants identified in *ARPP21.*** Analyses were performed with the ALS Knowledge Portal (ALSKP; individuals with ALS = 3864, controls = 7839) and Project MinE ALS sequencing consortium (individuals with ALS = 6596, controls = 2454) datasets. Variants only identified in controls were excluded from the figure (n=39). Seven missense variants from four peer-reviewed manuscripts are also indicated that were curated by the ClinGen Gene Curation Expert Panel (GCEP) in the review of *ARPP21*, none of which were predicted to be pathogenic by AlphaMissense.

**Supplementary Table 1.**
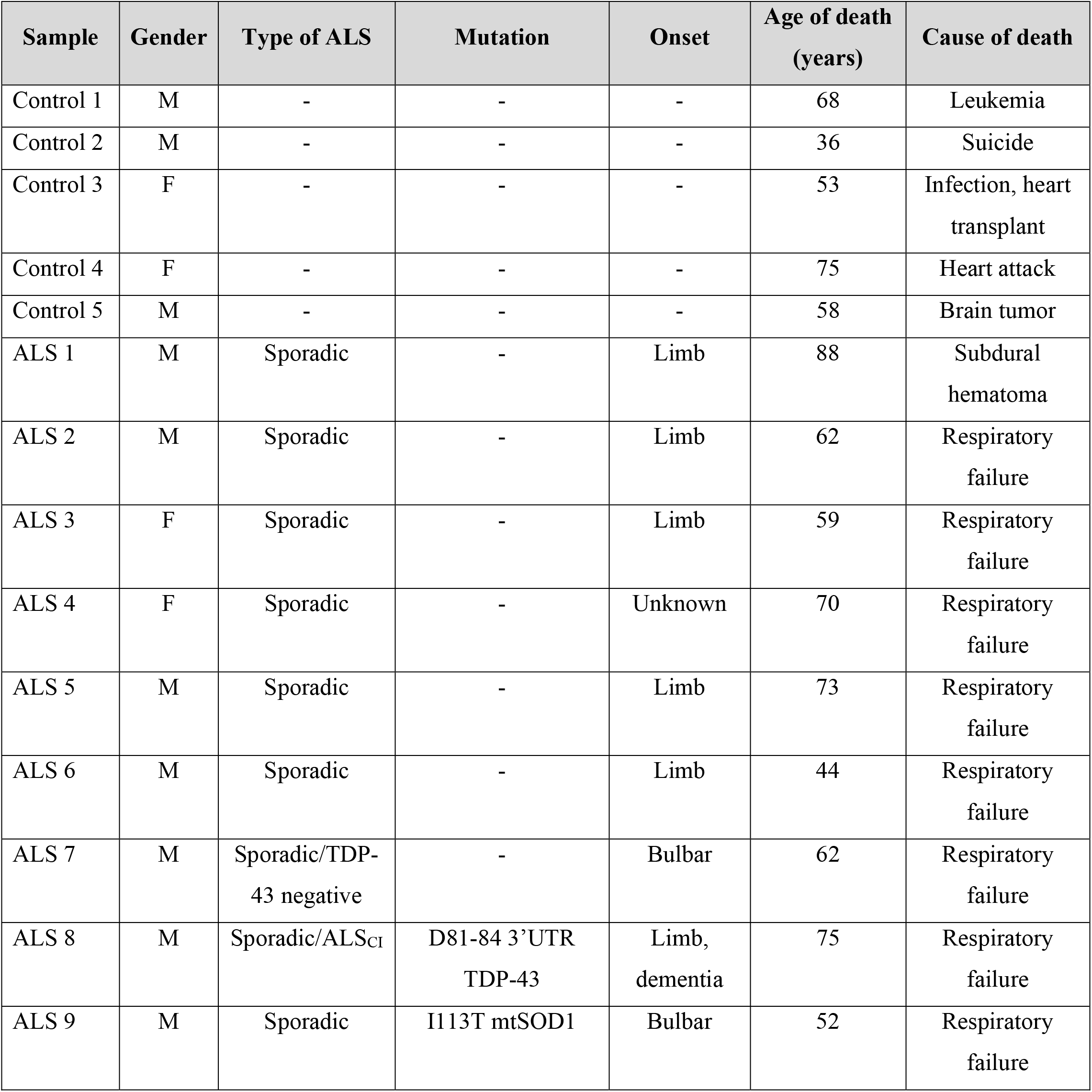
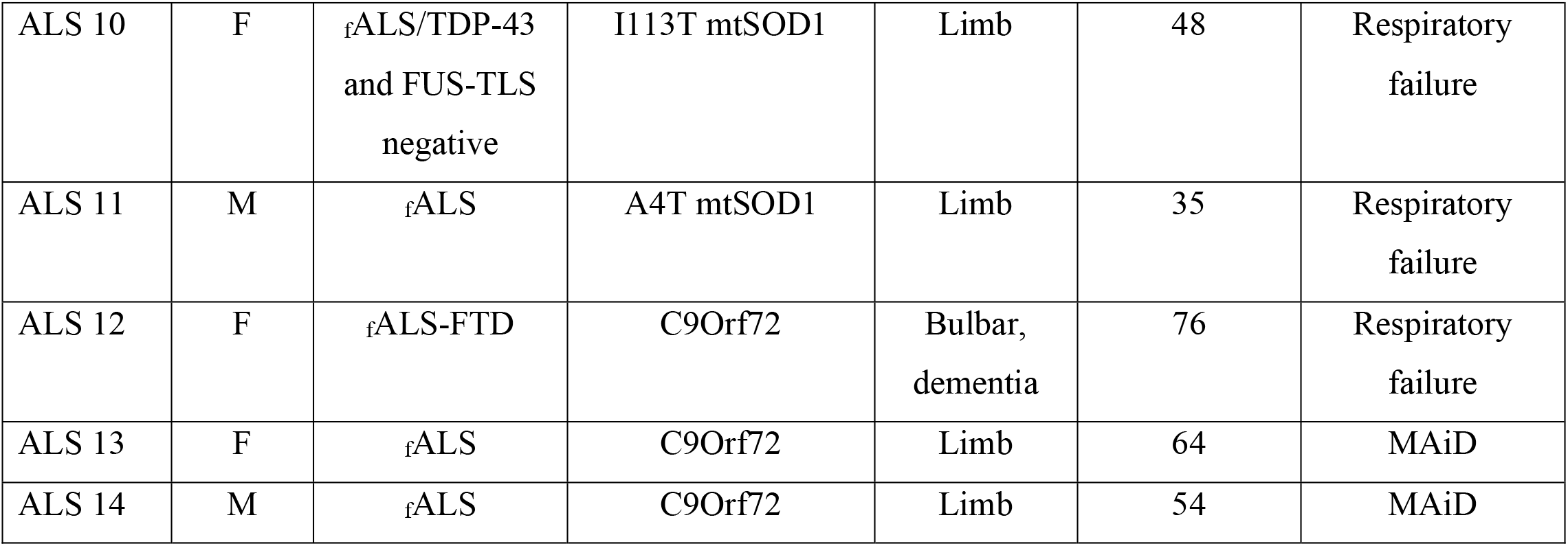
Patient demographics

**Supplementary Table 2.** Primary and secondary antibodies

| Antibody | Species | Company | Cat # | Dilution |
| --- | --- | --- | --- | --- |
| ARPP21 | mouse | Santa Cruz | sc-515480 | 1:100 |
| ARPP21 | rabbit | Proteintech | 11829-1-AP | 1:100 |
| F-actin | mouse | Abcam | ab130935 | 1:100 |
| flag | goat | Novusbio | NB600344 | 1:200 |
| G3BP | mouse | Abcam | ab56574 | 1:100 |
| GFP | goat | Abcam | ab6673 | 1:400 |
| myc | mouse | Cedarlane | CLX229AP | 1:250 |
| SZRD1 | rabbit | Proteintech | 24844-1-AP | 1:100 |
| TDP-43 | rabbit | Proteintech | 10782-2-AP | 1:500 |
| TDP-43 | mouse | Abcam | ab104223 | 1:500 |
| Ubiquitin | mouse | Millipore | MAB1510 | 1:500 |
| Anti-goat 488 | donkey | ThermoFisher | A-11055 | 1:1000 |
| Anti-mouse 488 | donkey | ThermoFisher | A-21202 | 1:1000 |
| Anti-rabbit 488 | donkey | ThermoFisher | A-21206 | 1:1000 |
| Anti-mouse 555 | donkey | ThermoFisher | A-31570 | 1:1000 |

**Supplementary Table 3.**
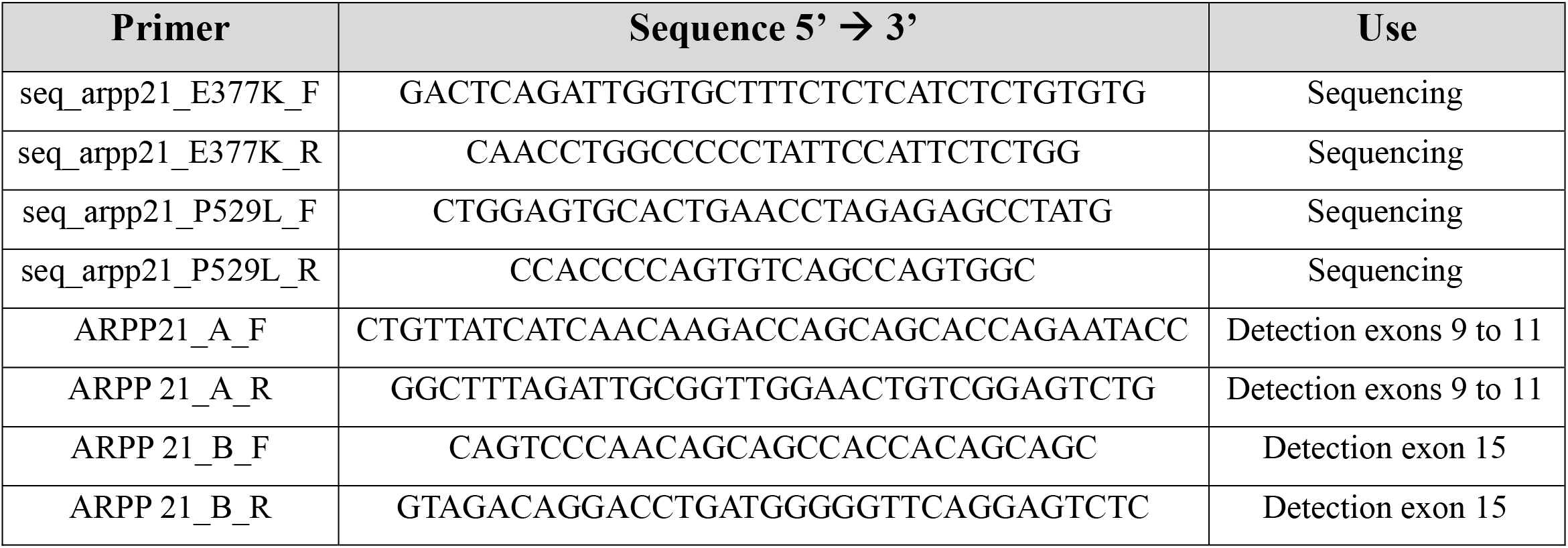
Primers

**Supplementary Table 4.** Plasmids

| Plasmid | Use |
| --- | --- |
| pcDNA-ARPP21<sup>wt</sup>-myc (Backbone: pcDNA3.1) | To express ARPP21<sup>wt</sup> tagged with myc at C-terminal |
| pcDNA-ARPP21<sup>E377K</sup>-myc (Backbone: pcDNA3.1) | To express ARPP21<sup>E377K</sup> tagged with myc at C-terminal |
| pcDNA-ARPP21<sup>P529L</sup>-myc (Backbone: pcDNA3.1) | To express ARPP21<sup>P529L</sup> tagged with myc at C-terminal |
| pcDNA-flag-SZRD1 (Backbone: pcDNA3.1) | To express SZRD1 tagged with flag at N-terminal |
| pHA-Clover (Addgene # 163366) | Backbone – Use to express clover alone |
| pTDP-43-wt - clover | To express TDP-43-wt tagged with clover at C-terminal |
| pARPP21<sup>wt</sup>-clover | To express ARPP21<sup>wt</sup> tagged with clover at C-terminal |
| pARPP21<sup>E377K</sup>-clover | To express ARPP21<sup>E377K</sup> tagged with clover at C-terminal |
| pARPP21<sup>P529L</sup>-clover | To express ARPP21<sup>P529L</sup> tagged with clover at C-terminal |
| pAAV-Syn-GFP (Backbone: pUC18) | To generate AAV9-GFP |
| pAAV-Syn-ARPP21<sup>wt</sup> (Backbone: pUC18) | To generate AAV9-ARPP21<sup>wt</sup> |
| pAAV-Syn-ARPP21<sup>P529L</sup> (Backbone: pUC18) | To generate AAV9-ARPP21<sup>P529L</sup> |

